# *Mycena* species can be opportunist-generalist plant root invaders

**DOI:** 10.1101/2021.03.23.436563

**Authors:** Christoffer Bugge Harder, Emily Hesling, Synnøve S. Botnen, Bálint Dima, Tea von Bonsdorff-Salminen, Tuula Niskanen, Susan G. Jarvis, Kelsey E. Lorberau, Andrew Ouimette, Alison Hester, Erik A. Hobbie, Andy F.S. Taylor, Håvard Kauserud

**Author notes:** Corresponding author: Christoffer B: Harder.

## Abstract

Recently, several saprotrophic genera have been found to invade/interact with plant roots in laboratory growth experiments, and this coincides with reports of abundant saprotrophic fungal sequences in plant roots. However, it is uncertain if this reflects field phenomena, and if reports on coincidentally amplified saprotrophs are simply coincidental.

We investigated root invasion by presumed saprotrophic fungi by focusing on the large genus *Mycena* in **1)** a systematic analysis of the occurrence of saprotrophic fungi in new and previously published ITS1/ITS2 datasets generated from roots of 10 mycorrhizal plant species, and **2)** we analysed natural abundances of ^13^C/^15^N stable isotope signatures of fungal/plant communities from five comparable field locations to examine the trophic status of *Mycena* species.

*Mycena* was the only saprotrophic genus consistently found in 9 of 10 plant host roots, with high within-host variation in *Mycena* sequence proportions (0-80%) recovered. *Mycena* carpophores displayed isotopic signatures consistent with published ^13^C/^15^N profiles of both saprotrophic or mutualistic lifestyles, with considerable intraspecific variation, resembling the patterns seen in growth experiments. These results indicate that multiple *Mycena* species opportunistically invade the roots of a range of plant species, possibly forming a spectrum of interactions. This potentially challenges our general understanding of fungal ecology.

**Originality significance statement:** This is the first study to apply a dual approach of systematic metabarcoding of plant roots and stable isotope signatures on dried field material to the large and common saprotrophic fungal genus *Mycena*. This is significant as it shows that members of this genus, normally not expected to be found inside plant roots at all, are in fact associated eith multiple plant hosts. The study furthermore shows that species in this genus may occupy different ecological roles in the field besides being saprotrophic. That a large and common fungal genus known to be a quantitatively important litter decayer can be an opportunistic root invader and interact with host plants is of interest to all mycologists and ecologists working on plant-fungus/microb symbiosis.

## Introduction

Among ecologists, a consensus is emerging that the classical assignment of fungal species into single trophic groups with a mycorrhizal, saprotrophic or pathogenic lifestyle may be too restrictive (Baldrian and Kohout, 2017; Selosse *et al*., 2018). Some otherwise free-living fungi have been shown to invade plant roots and exist as either asymptomatic endophytes (neither harmful nor beneficial), and then switch from endophytic into becoming pathogenic (e.g. *Fusarium graminearum*, Lofgren *et al*., 2018), or into forming AM mycorrhizas (*Piriformospora indica*, Weiß *et al*., 2016) or ericoid mycorrhizal associations (*Meliniomyces* spp., Martino *et al*., 2018).

A wide range of saprotrophic fungal genera have been screened for their ability to colonise *Pinus sylvestris* and *Picea abies* seedling roots *in vitro* (Smith *et al*., 2017), and several, including *Mycena, Gymnopus, Phlebiopsis, Marasmius* or *Pleurotus*, invaded roots apparently without decomposing dead tissue in the process. However, beyond the invasion, the nature of the interactions with the plant host (if any) remains unknown.

For *Mycena*, however, there are several lines of direct and indirect evidence for their ability to invade and interact with living plant roots, at least *in vitro. Mycena* species have been identified as potential orchid mycorrhizal symbionts (Ogura-Tsujita *et al*., 2009; Zhang *et al*., 2012), endophytes in photosynthetic moss tissue (Davey *et al*., 2013) and non-mycorrhizal brassicaceous plants (Glynou *et al*., 2018). They have also been shown to form mycorrhiza-like structures in roots of *Vaccinium corymbosum* in growth studies (Grelet *et al*., 2017). Recently and perhaps most importantly, Thoen *et al*. (2020) showed that multiple species and individual strains of *Mycena* could colonise roots of *Betula pendula* seedlings in vitro, and formed a gradient of interactions from harmful to neutral to beneficial, with some species/strains being able to transfer nutrients to the plant host. This is significant, as prior to this *Mycena* sensu stricto (Moncalvo *et al*., 2002) (henceforth simply “*Mycena*”), which is one the largest genera in Agaricales (over 500 species), widespread across habitats and climate zones (Kühner, 1938; Maas Geesteranus, 1992; Rexer, 1994; Robich, 2003; Aronsen and Læssøe,2016), was known primarily as quantitatively important litter and wood debris decomposers (Boberg *et al*., 2008; Baldrian *et al*., 2012; Kyaschenko *et al*., 2017). The spectrum of interactions seen in Thoen *et al*. (2020) is further noteworthy in the light of the “Waiting room hypothesis” (van der Heijden *et al*. 2015) on mycorrhizal evolution, which suggests that the mycorrhizal habit evolves from saprotrophs gradually via neutral endophytic intermediate states. This hypothesis has remained controversial even though it is accepted that the mycorrhizal habit has evolved on numerous, independent occasions from saprotrophic ancestors (Tedersoo and Smith 2013, Kohler *et al*. 2015). Thus, the genus *Mycena* may represent a promising research model for studying both ecological versatility in fungi and the possible ongoing evolution of fungi traditionally believed to be purely saprotrophic *en route* to developing mycorrhizal abilities.

Most of this evidence for trophic versatility in *Mycena* originates from *in vitro* studies, and it is uncertain to what extent this translates to the field. To investigate the trophic mode of fungi in natural environments, analysis of the natural abundance of ^13^C:^12^C and ^15^N:^14^N ratios (isotope ratios, expressed as d^13^C and d^15^N values relative to known standards) can be applied directly to fungal carpophores and other field material. Mycorrhizal fungi are generally more enriched in ^15^N and depleted in ^13^C than saprotrophic fungi (Taylor *et al*., 1997; Kohzu *et al*., 1999; Hobbie *et al*., 1999; Hobbie *et al*., 2001; Griffith, 2004; Mayor *et al*., 2009). Based on comparisons between fungi of known trophic status, natural abundance of 15 and 13C can also give strong indications of the nutritional mode of fungi with unknown trophic status. Thus, Halbwachs *et al*. (2018) recently used this approach to strongly suggest that *Hygrocybe*, another genus traditionally believed to be saprotrophic, was most likely biotrophic with plants. The occurrence and abundance of *Mycena* sequences retrieved from wild plant roots also suggests interactions with plant roots in situ. There are scattered reports of *Mycena* identified from inside wild plant roots, particularly in the Arctic plants, including *Bistorta vivipara, Cassiope tetragona, Dryas octopetala*, and *Salix polaris* (Blaalid *et al*., 2014; Botnen *et al*., 2014; Lorberau *et al*., 2017). In some cases*, Mycena* sequences comprised >30-50% of the total reads, suggesting that they are not simply casual colonisers. It has been suggested that the harsh and oligotrophic Arctic environments stimulate otherwise free-living fungal genera (including *Mycena*) to explore new ecological niches (Jumpponen and Trappe, 1998; Ryberg *et al*., 2009; Ryberg *et al*., 2011; Timling *et al*., 2012; Botnen *et al*., 2014; Lorberau *et al*., 2017). Nevertheless, *Mycena* reads have also been recovered in high quantities from inside living *Picea abies* roots in temperate environments (Kohout *et al*., 2018). In this case, *Mycena* species were present in the roots prior to felling but then became dominant post-felling. Overall, however, current information on the occurrence of *Mycena* and other saprotrophs in roots unsystematic and too scattered to identify any clear patterns of their occurrence and abundance.

High throughput sequencing (HTS) studies of fungal communities in plant roots are generally targeting mycorrhizal fungi (Buee *et al*., 2009; Tedersoo *et al*., 2010; Bahram *et al*., 2011; Blaalid *et al*., 2014; Vasar *et al*., 2017; Kaur *et al*., 2019), and the workflow requires the annotation of tens of thousands of OTUs/clusters of fungi into ecological guilds. Most studies tend to favour more conservative taxonomic ecological generalisations based at the genus level, which is traditionally considered the most relevant level for separating fungal taxa by nutritional mode (Fries and Mueller, 1984; Molina and Trappe, 1994; den Bakker *et al*., 2004; Tedersoo and Smith, 2013; Garnica *et al*., 2016). Ecological annotation software (Nguyen *et al*., 2016) is largely based on this view. Thus, in studies of mycorrhizal fungi in roots, fungal genera identified as being saprotrophic may be at best briefly mentioned and/or reported as one lumped ecological group (Menkis *et al*., 2012; Tedersoo and Smith, 2013), or simply dismissed as accidental contamination (Liao *et al*., 2014). This means that the ecological understanding of root ecosystems may become oversimplified, and large quantities of potentially informative data are left unanalysed and erroneously ignored.

Here, we present a dual analysis on the occurrence of *Mycena* in roots of wild plants in a range of ecosystems, and an investigation of the potential trophic versatility of Mycena in the field by presenting: 1) a systematic analysis of data from 10 plant species from Arctic and temperate regions (Information on plants and studies given in Table 1) from previously published and newly generated ITS1/ITS2 HTS data sets from living plant roots, and 2) a comparison of the natural abundance of ^13^C and ^15^N in carpophores of *Mycena* with other fungi, soils and mycorrhizal host plants from 253 fungal collections, host plants and soils from five field locations (See Fig S1-S2 for a map of sampling sites).

**Table 1.** Overview of all species and samples included in this study. ^1^Number represent samples which ended up being included in the final analyses. Our plant samples consisted of the herbaceous ectomycorrhizal Arctic *Bistorta vivipara*, subshrub *Dryas octopetala*, and the dwarf shrub *Salix polaris* (Blaalid et al., 2012; Yao et al., 2013; Blaalid et al., 2014; Botnen et al., 2014; Davey et al., 2015; Mundra et al., 2015); the Arctic ericaceous *Cassiope tetragona* (Lorberau et al., 2017) and the ectomycorrhizal conifer *Pinus sylvestris* (Jarvis et al., 2015) from Scotland; and new data also from Scottish (temperate) plants: the arbutoid mycorrhizal *Arctostaphylos alpina* and *A. uva-ursi*, ectomycorrhizal dwarf shrubs *Betula nana* and *Salix herbacea*, and the ectomycorrhizal trees *Betula pubescens* and additional *Pinus sylvestris*. The *Betula pubescens* samples all came from saplings of < 1 m kept low by deer/sheep grazing; the other host plants collected were mature.

| ITS1<br>Title of study | Samples<sup>1</sup> | Locality | Description of original purpose |
| --- | --- | --- | --- |
| Blaalid et al. (2012) | *Bistorta vivipara* (n=59) | Finse(Nor way) | Succession |
| Yao et al. (2013) | *Bistorta vivipara* (n=51) | Finse(Nor way) | Ridge-to-snowbed gradient |
| Blaalid et al. (2014) | *Bistorta vivipara* (n=146) | 32 localities on Svalbard | Spatial/bioclimate variation |
| Botnen et al. (2014) | *Bistorta vivipara* (n=19), *Dryas octopetala* (n=22), *Salix polaris* (n=20) | Blomsterdalen, Svalbard | Host specificity |
| Mundra et al. (2015) | *Bistorta vivipara* (n=84) | Isdammen, Svalbard | Temporal variation |
| Davey et al (2015) | *Bistorta vivipara* (n=103) | Svalbard, Finse(Nor way) | Succession/glacial chronosequence gradient |
| Botnen et al. (2019) | *Bistorta vivipara*(n=57) | Scotland, Austria, Iceland, Jan Mayen | Biogeography |
| ITS2<br>Title of study |  |  |  |
| Jarvis et al. (2015) | *Pinus sylvestris* (n=32) | Scotland | Altitude |
| Lorberau et al (2017) | *Cassiope tetragona* (n=15) | Endalen, Svalbard | Drought |
| Biogeography project (new data) | *Arctostaphylos alpine*(n=10), *Arctostaphylos uva-ursi*(n=8) *Betula nana* (n=8) *Salix herbacea* (n=7) | Scotland | Biogeography |
| Altitude project (new data) | *Arctostaphylos uva-ursi*(n=68), *Pinus sylvestris* (n=9) | Scotland | Altitude |
| Birch (new data) | *Betula pendula* (n=81) | Scotland | Grazing effects |

We investigated four main research questions: **1)** Are *Mycena* species (and other supposedly saprotrophic genera) systematically found inside living plant roots in significant quantities? **2)** If so, are these more prevalent in Arctic/Alpine environments? **3)** Are there indications of host preferences/specificity among invasive *Mycena* species? **4)** Do the data from ^13^C and ^15^N abundances support the view that *Mycena* species may form mutualist association with plants?

## Results

### HTP-sequencing data summary

Our dataset consisted of 3 species amplified with ITS1 primers and 2 species amplified with ITS2 primers from previously published data reanalysed here. 5 new species (from three separate 454 datasets amplified with ITS2 primers) represent new data (Table 1).

After quality sorting, a final dataset of 889,290 ITS1 sequences were clustered into 1193 3% OTUs (n≥1 0, henceforth simply OTUs) for the three plant species where the ITS1 marker were used, and an ITS2 dataset of 992890 sequences and1032 3% OTUs in the seven datasets with ITS2 (Table 2). For a detailed list of the sequence sorting steps on the ITS1 and ITS2 data and the respective counts for each host species, see Supplementary data and Tables S1-6. Applying the “coverage/completeness” method for assessing saturation (Chao and Jost, 2012), 111 samples failed to meet the 97% coverage cutoff value and were discarded. Though the iNEXT (Hsieh *et al*., 2016) extrapolations of observed species richness suggested that some slight undersampling remained in some samples (Fig S5), none of the ten species showed correlations between *Mycena* infection levels (all Ps>0.05, table S7) with sampling depth. We analysed the datasets with both amplicon sequence variants (ASVs/”zotus”, aimed at capturing the haplotype diversity) and 3%-OTUs, aimed at capturing species. As expected, there were higher numbers of ASVs than OTUs. However, using 3% OTUs or ASVs made little difference to the taxonomic composition of the datasets, as seen by the near-identical *Mycena* shares in OTUs and ASVs. There were no clear signs of potential host specialisation by *Mycena* at either the finer ASV-(“haplotype”) scale or at the coarser 3% OTU scale (Tables S8-9). Thus, the further analyses focused on the 3% OTU datasets.

**Table 2.** Total sequence counts for OTU/ASVs for each host species.^2^Numbers represent samples included in the final analyses.

|  |  |  |  |  |  |  |  |
| --- | --- | --- | --- | --- | --- | --- | --- |
| ITS1 | B. vivipara (n=519)<sup>1</sup> |  | D. octopetala (n=22) |  | S. polaris (n=20)<sup>2</sup> |  |  |
| OTUs (n=1193) | 803649 |  | 39880 |  | 30069 |  |  |
| ASVs (n=2272) | 908621 |  | 43286 |  | 33141 |  |  |
| ITS2 | A.alpine (n=10) | A.uva-ursi (n=76)<sup>1</sup> | B.nana (n=8) | B.pendula (n=81)<sup>2</sup> | P. sylvestris (n=41) | S. herbacea (n=7) | C. tetragona (n=15) |
| OTUs (n=1032) | 59829 | 241418 | 49927 | 90214 | 212577 | 25298 | 313627 |
| ASVs (n=1559) | 61703 | 247091 | 51182 | 93429 | 219124 | 25497 | 385763 |

For ITS1, 606 of 1193 OTUs (78.3% of the sequences) could be identified to genus level by SINTAX at the threshold of BPP >0.6; for ITS2, this number was 513 of 1032 ITS2 OTUs (84.5% of sequences).

The same SINTAX classification identified 13 *Mycena* OTUs (1.5% of all ITS1 sequences) in the ITS1 dataset, and 14 *Mycena* OTUs in ITS2 (12.6% of all ITS2 sequences). However, in a second identification step especially targeting *Mycena* where representative sequences of all OTUs were clustered with the *Mycena* ITS database 576 sequences (described below), an additional 7 ITS1 OTUs and 7 ITS2 OTUs not identified as *Mycena* by SINTAX at BPP >0.6 formed clusters (at 97%) with *Mycena* species in the database. Thus, in total 20 ITS1 OTUs (2.1% of sequences) and 21 ITS2 OTUs (15.8% of sequences) could be identified as *Mycena* (s. str) with these two combined methods. (3 ITS1 OTUs identified as *Mycena* by the 8.2 utax eukaryote reference database represented taxa now classified as *Phloeomana* and *Atheniella* (Redhead, 2013) and were excluded from detailed analysis).

Of other (*non-Mycena*) taxa traditionally considered to be saprotrophic/endophytic, we found Sebacinales in the four Arctic host plants, the zygomycete *Mortierella* in most Scottish hosts, and *Phialocephala in B. vivipara* and *Clavulinopsis/Clavaria* in *C. tetragona* (Fig. 2a-c). No other saprotrophic/endophytic genera formed more than 0.5% of the sequences in any of the host plants. The high and very variable infection levels and frequency patterns of *Mycena* were not found in any other saprotrophic/endophytic taxa.

**Fig.1.**
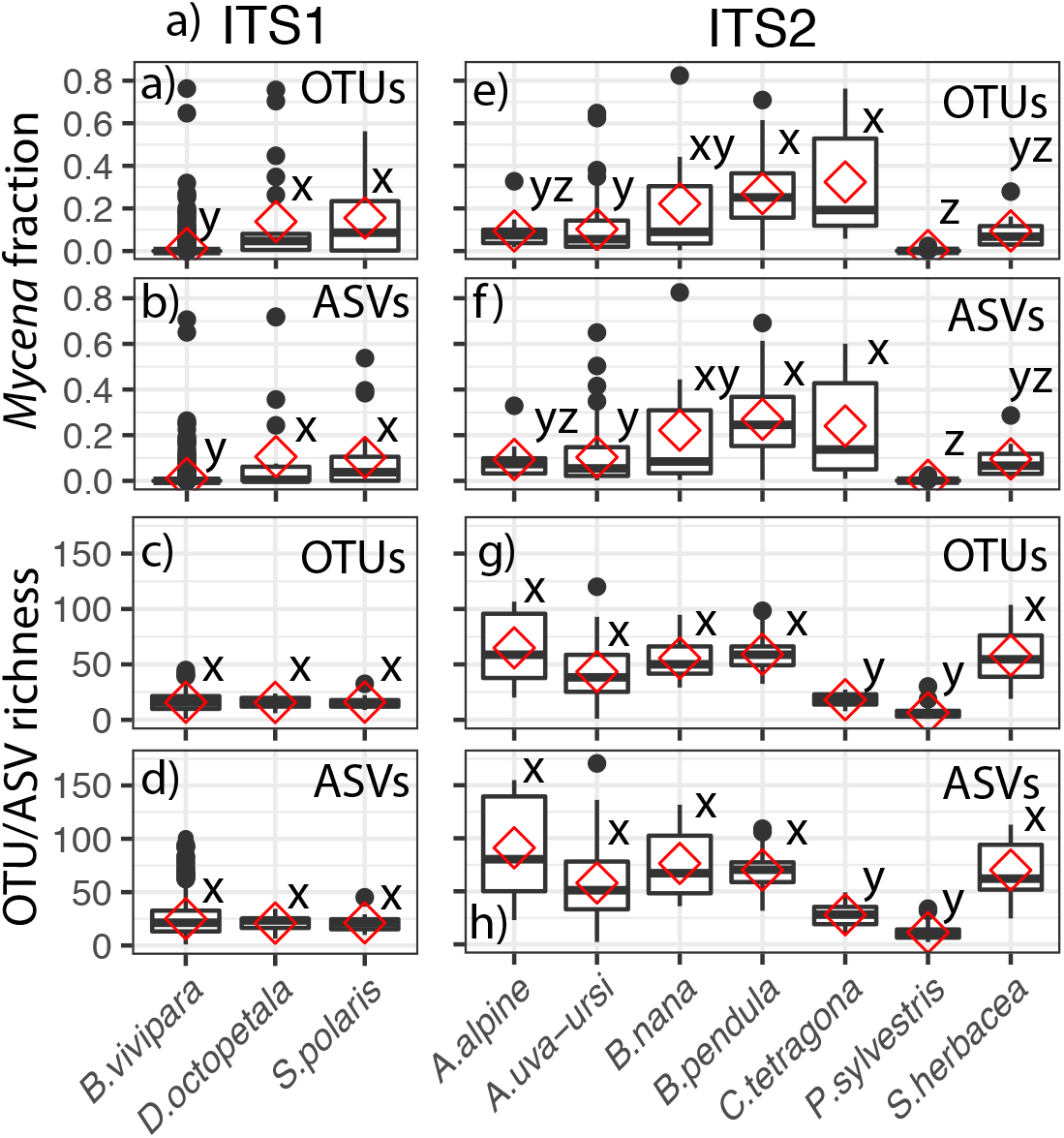
*Mycena* infection levels (fraction of read shares) and species richness at 97% coverage corrected for sampling depths (Chao and Jost 2012) for the ITS1 (a-d) and ITS2 (e-h) datasets. Very little differerence between the OTU and the ASV approaches were found. Letters x-y-z denote host species «significance groups» as found by ANOVA + Scheffes multiple comparisons test for a significance at the P< 0.05 level. Species sharing one identical letter (x, y or z) do not significantly differ from each other in mean *Mycena* infection level/overall species richness (at 97% coverage).

**Fig.2.**
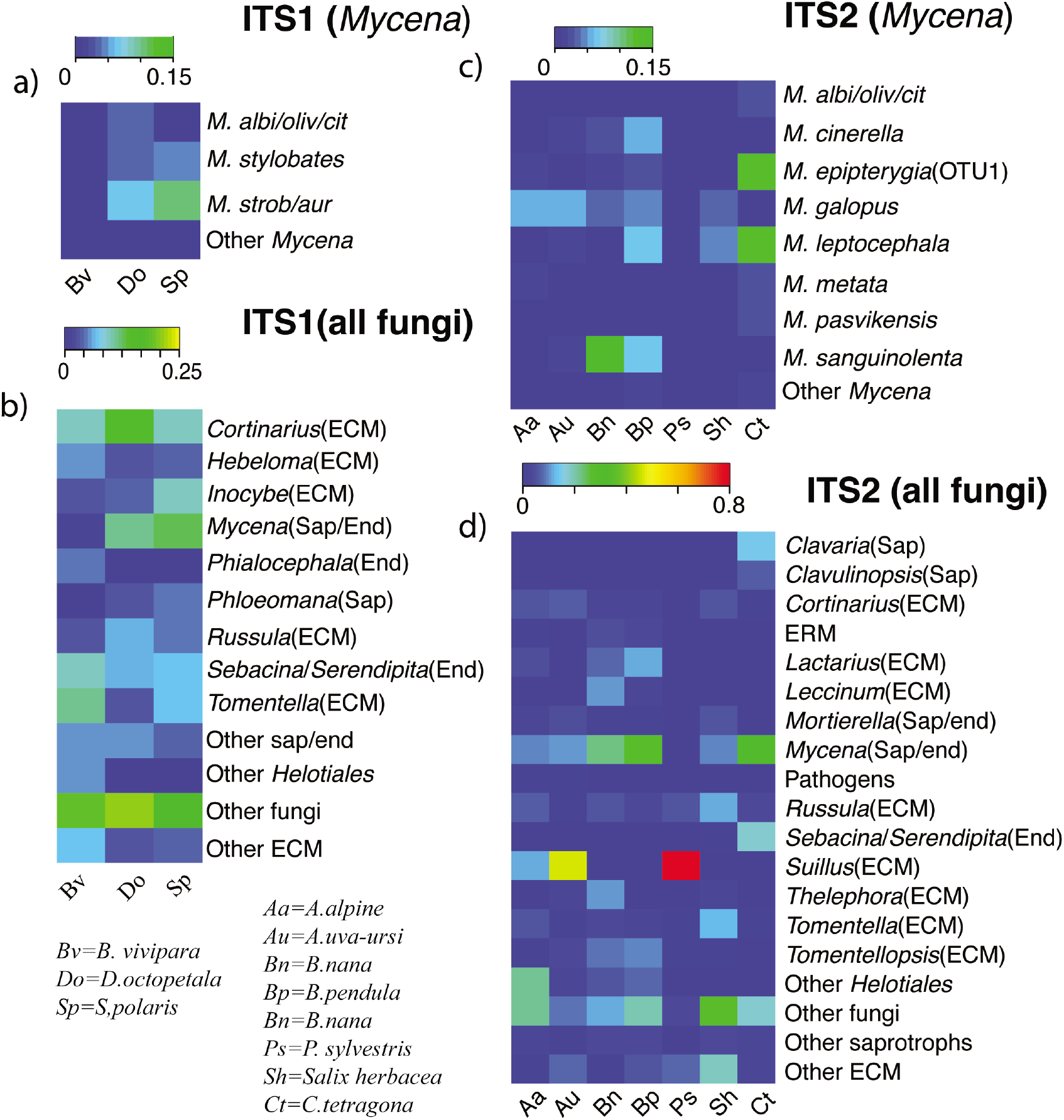
Heatmap of *Mycena* species occurrence (a-c) and overall genus occurrence (b-d). Note the slightly different colour bars in each plot. Only *Mycena* species that made up >1% of at least one sample were included as separate rows in a-c. In b-d, only genera that comprised >5% of reads as an average in at least one host species had its own separate row. Besides these criteria, we also included Helotiales that could not be identified to generic level, but might still conceivably harbour ericoid mycorrhizal fungi or dark septate endophytes (as *Phialocephala*). but might. ERM=Ericoid mycorrhiza, ECM= Ectomycorrhiza, Sap=Saprotroph, End=Endophyte. Note the near-complete dominance of ectomycorrhiza (particularly *Suillus*) in *P. sylvestris*. (Ps).

### *Mycena* infection levels

In nine out of ten host plants, *Mycena* infection levels reached 25-80% of all reads in individual samples (Fig. 2a-c), with considerable intraspecific infection variation (Fig 1, Fig. S3). For the ITS1 data, *Mycena* average read content for all 519 *B. vivipara* samples was significantly lower (than for *S. polaris* (n=20) and *D. octopetala* (n=22) (Fig.1a-b, Table S10)). However, the *S. polaris* and *D. octopetala* data sets came from only one locality (Botnen *et al*., 2014), and when comparing them only with the *B. vivipara* dataset (n=19) from the same locality, no significant differences were observed (1-way ANOVA, df=2, F=1.36, p=0.263). Without considering the differences in sample sizes (Table 1), ITS2 host species could be roughly divided into three groups based on average *Mycena* infection level - 1) *P. sylvestris* with virtually no *Mycena*, 2) an intermediate group (median values about 5-10%) with *S. herbacea, A. alpine, B. nana*, and *A. uva-ursi*, and 3) *B. pubescens* and *C. tetragona* with median *Mycena* infection levels above 20%. While all species (except *P. sylvestris*) harboured individual samples with few if any Mycena and some with >30%, there were still significant differences between these three rough categories.

(Fig. 1 d-e, Table S11)).

### Environmental influences on Mycena

Disparities in sample size and study metadata only permitted limited testing of environmental influences on *Mycena* infection to three host species. In *C. tetragona*, there were no difference in *Mycena* infection levels between samples derived from drought and control plots applied by Lorberau *et al*. (2017)(two-tailed *t*-test, unequal variances, p=0.57). In *A. uva-ursi*, the level of *Mycena* infection decreased significantly with increasing altitude (65-805 m above sea level) (R^2^=0.2, p<0.0001, data not shown), which is contrary to the assumption (question 2) that increasingly stressful environments facilitating infection with *Mycena*/saprotrophs.

In *B. vivipara*, no correlations between *Mycena* infection level and annual mean temperature, latitude nor mean temperature of the wettest quartal were found (all adjusted R^2^<0.01, all P>0.25, see Table S12). A very weak correlation between decreasing *Mycena* species richness in roots and increasing mean temperature of the wettest quartal (R^2^=0.01, P=0.04) disappeared when the Austrian outlier samples (which contained no *Mycena*) were excluded.

A chi-quare test on the observed vs. expected prevalence of *Mycena* in 222 *B. vivipara* host plants from 44 patches (a patch constituted multiple plants collected in close proximity) showed a significant, non-random association (c^2^=65.22, df=43, p=0.01) of *Mycena* infections in host plants, suggesting that *Mycena*-infected *B. vivipara* were distributed in clumps.

### *Mycena* phylogenetics of OTUs and species diversity

Among the 20 ITS1 and the 21 ITS2 OTUs that were identified as *Mycena*, we found no phylogenetic signal suggesting that root invasion might be linked to certain clades (Fig. 3). Many of the same *Mycena* species were found in both the ITS1 and ITS2 datasets with 12 ITS1-ITS2 pairs of OTUs clustered with >90% probability to the same branches. There were no indications of host specialisation by *Mycena* species on particular host species, with large individual variations in all host plants (again except *P. sylvestris*) between which *Mycena* species that were found (Fig. S3), Several *Mycena* species such as *M. epipterygia* or *M. leptocephala* occurred in infection levels of >10% in at least one individual of 6 of 10 host species or more (Fig. S3). Indeed, the two OTUs were the only OTUs shared between *C. tetragona* from Svalbard and all Scottish host species (except *P. sylvestris*).

**Fig.3.**
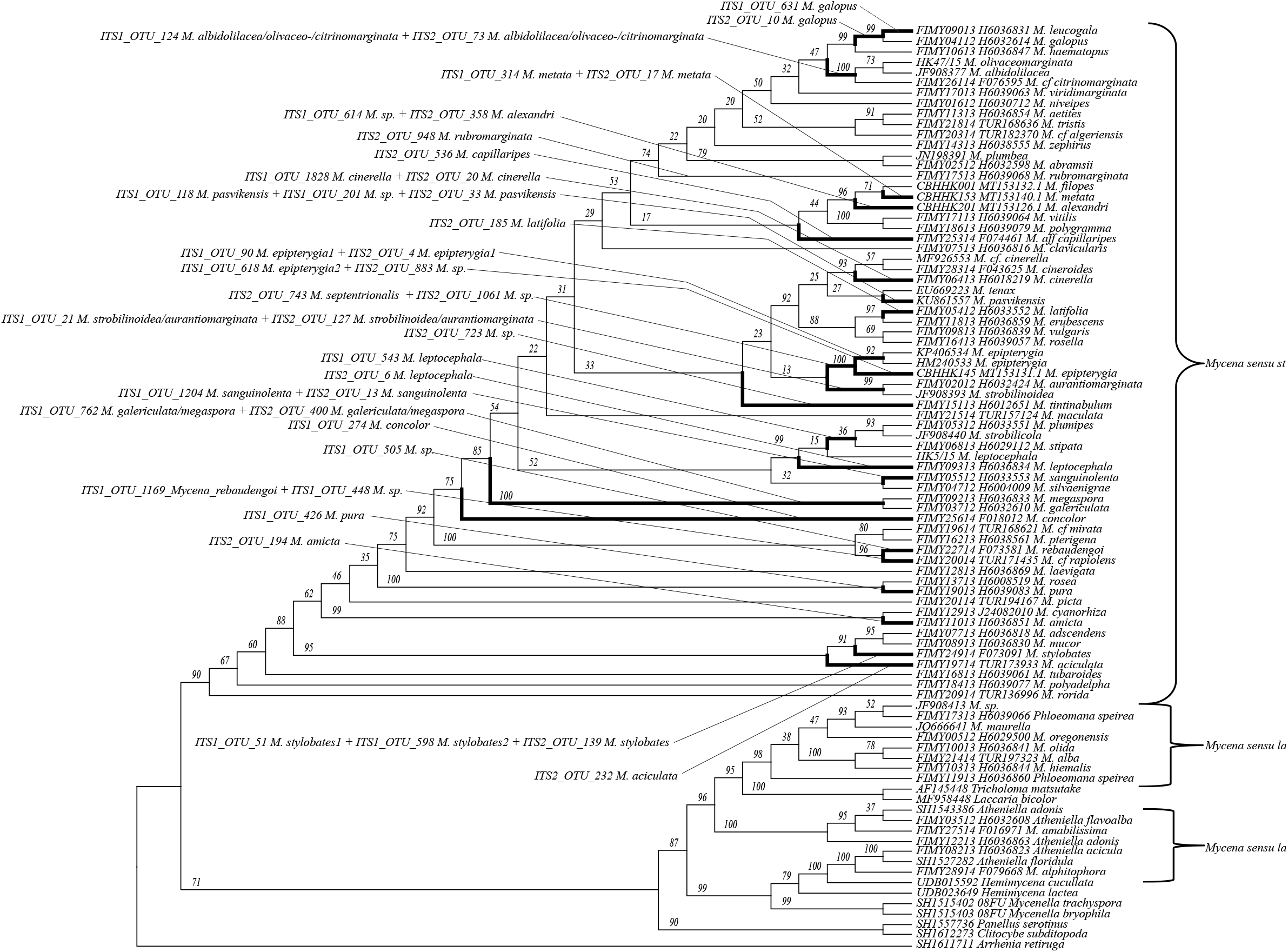
RaxML complete ITS phylogeny of 64 *Mycena* s.s. sequences, with 25 outgroups from “*Mycena* s.l.” and other Agaricales. Bootstrap supports indicated about each branch. 20+21 OTUs from ITS1 and ITS2 are superimposed upon the branches with the best fit.

### Mycena database

We compiled 576 new and previously published full-length Mycena ITS Sanger sequences representing 137 identified species level into a database (see Experimental procedures below).They clustered into 156 and 139 ITS1 and ITS2 3% OTUs, respectively.

For both regions, some OTUs contained two or more species (such as *Mycena galericulata + M. megaspora* and *M. olivaceomarginata, M.citrinomarginata* and *M. albidolilacea*), while other species were split into multiple OTUs (e.g. *M. pura* and *M. epipterygia)*. Average intraspecific variation was 3.6% (ITS1) and 2.7% (ITS2) (Fig. S4).

### Stable isotope data

On average, carpophores values of 15N and 13C placed *Mycena* among the saprotrophs. They were higher in δ^13^C and lower in δ^15^N than the average of the remaining *non-Mycena* saprotrophs (Fig. 4a-e). The δ^13^C values of all saprotrophic species for all regions were between −26 and −22‰, except for a *Phloeomana speirea* at Finse (Fig. 4a) at −27.1‰, and one *Mycena metata* collection from Svalbard (Fig. 4c) at −26.9‰. However, there were striking anomalies (and intraspecific variations) in the δ^15^N values for certain individual collections of *Mycena*, particularly *M. pura*, which varied between 1.7‰ for *M. pura* in Gribskov to 12.6‰ for *M. pura1* at Finse. A *t*-test showed the *M. pura1* and *M. pura2* collections at Finse to be strongly and significantly higher in δ^15^N than the average for the other *Mycena* at Finse (p < 0.0001), and slightly but still significantly higher in δ^13^C (p = 0.02).

**Fig.4.**
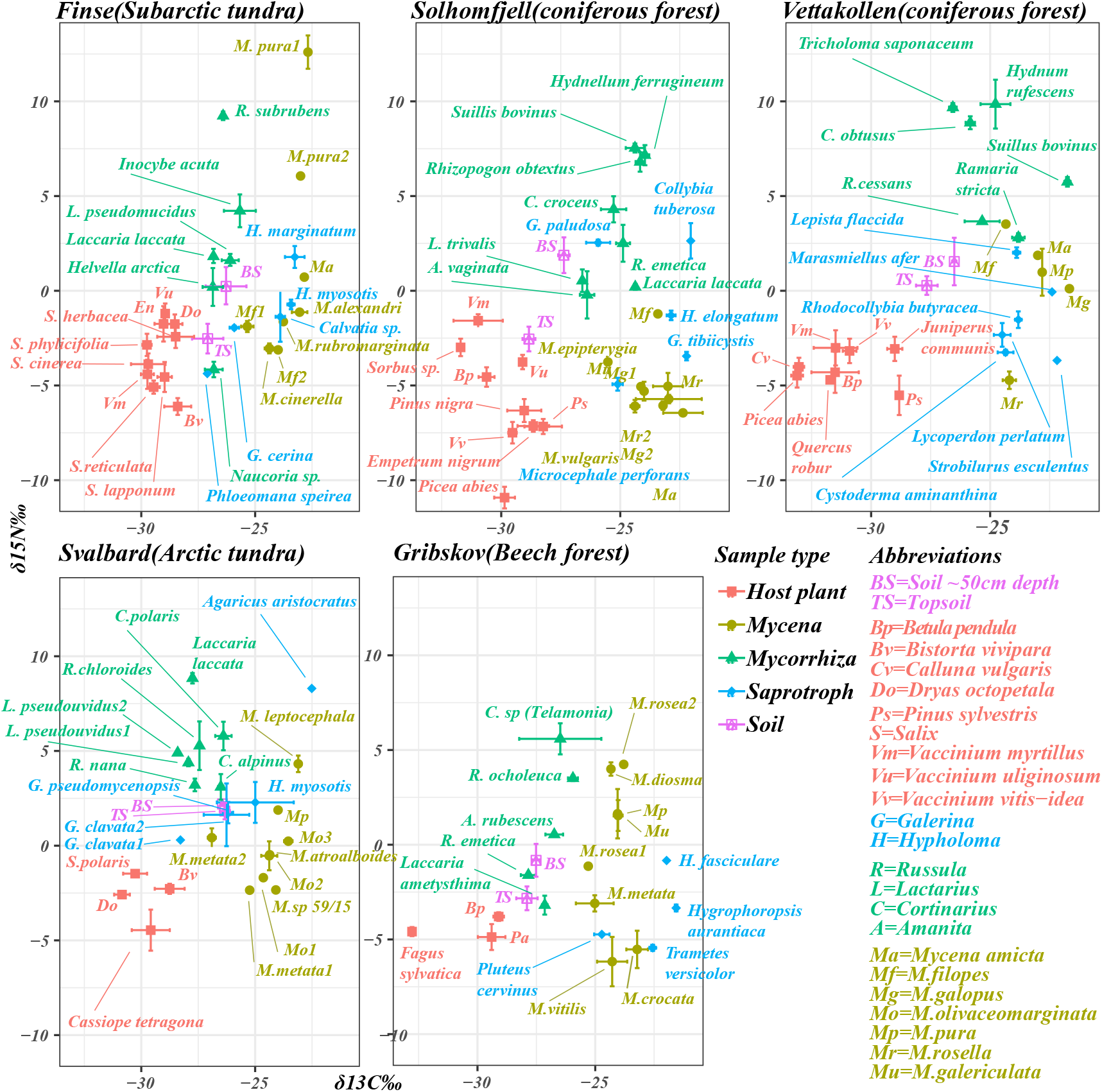
Biplots of stable isotopes ^15^N and ^13^C for soil, host plants, and the three simplified categories *Mycena*, Mycorrhiza and saprotrophs. Overall, saprotrophs will be found predominantly bottom right, mycorrhizal fungi top left. Note the deviant particular *M. puras* at Finse, upper left.

In the stepwise regression of δ^15^N, genera were separated by up to 12‰ into five groups (Table S13). Relative to the mean, *Mycena* grouped at −3‰ with the litter decay fungi *Calvatia, Lycoperdon*, and *Rhodocollybia*. The overall adjusted r^2^ of the regression model was 0.56, with site accounting for 8.1% of variance and the remaining 48.0% accounted for by genus.

In the stepwise regression of δ^13^C, genera were separated by up to 5‰ into eight groups (Table S14). Relative to the mean, *Mycena* grouped at +1‰ with both the ectomycorrhizal *Rhizopogon*, the partly saprotrophic/ectomycorrhizal *Ramaria* and the litter decay fungi *Calvatia, Lepista*, and *Rhodocollybia*. The overall adjusted r^2^ of the regression model was 0.66, with site accounting for 11.7% of variance, nitrogen concentration (%N) for 2.8%, and the remaining 51.4% accounted for by genus.

The δ^15^N values for host plants in all 5 regions were all below 0‰, which is significantly lower than not only carpophores, but also the soil (Fig.4). Soil δ^15^N values were below 2‰, with lower overall N contents but higher δ^15^N values for deeper soil depths, in line with earlier studies (Evans, 2007; Seitzman *et al*., 2011, Clemmensen K, 2013; Halbwachs *et al*., 2018). The δ^15^N and δ^13^C values differed significantly between the five regions, but the amount of variance for both measures explained by region in the mixed linear model was <20% for both isotopes and well below that explained by sample type and genus/sample type in combination (> 75% for both).

Overall, the other fungal genera had isotopic profiles that matched their expected nutritional mode.

## Discussion

This study constitutes the first systematic analysis of *Mycena* in wild plant roots, and the results clearly indicate that *Mycena* species are frequent root colonisers of a taxonomic range of mycorrhizal host plants, although infection levels are very variable. The isotopic data here suggest that they could have several potential ecological functions inside the roots. They could be endophytes, as has been suggested for Sebacinales (Blaalid *et al*., 2014; Botnen *et al*., 2014; Lorberau *et al*., 2017) or dark septate endophytes (Newsham, 2011) in many Arctic plants. However, here we found *Mycena* infection to be widespread in Arctic/alpine hosts as well as in temperate hosts, and the general lack of host-specificity in *Mycena* was also universal. The ability to colonise living roots appears to be a widely shared trait across the *Mycena* phylogeny, consistent with the findings of Thoen *et al*. (2020).

*Mycena* infections displayed a qualitatively remarkably similar pattern in 9 of 10 host plants: present in many samples and varying from little or no infection up to >40-50% of the recovered reads i.e:. The complete lack of *Mycena* (and saprotrophs in general) in *P. sylvestris* (Fig.1) is noteworthy, as *Mycena* were frequent root invaders in *P. sylvestris* seedlings in *in vitro* growth experiments by Smith *et al*. (2017). Pines have strongly heterorhizic root systems and only the final feeder roots are not either suberised or metacutinised, which will severely limit colonisation by fungi. In addition, under field conditions ectomycorrhizal colonisation levels of the feeder roots will be close to 100%, so the available surface for colonisation by non-mycorrhizal fungi will be very limited.

However, Kohout *et al*. (2018) found *Mycena* to be widespread in roots of mature (+80y) stands of conifers (*P. abies*) in forests. Furthermore, the *Mycena* infection patterns in *B. pendula* seedlings *in vitro* (Thoen *et al*., 2020) were indeed consistent with our observations in the close relative *B. pubescens* roots in the field.

However, the spruce in Kohout *et al*. (2018) originated from intensely managed and ultimately clear-cut forest, and they had a high overall root fungal diversity in addition to the levels of *Mycena* infections. The *B. pubescens* in this study were tree saplings of >1 m on a location where they were kept low by a particularly high grazing impact from sheep and deer, and all other species with high *Mycena* infection levels were smaller dwarf shrubs of varying sizes or herbaceous plants, also known to be subject to deer/reindeer grazing (Kolari *et al*., 2019). In contrast, the *P. sylvestris* in our study were in largely undisturbed stands within a national park, and their near-complete dominance by one genus (*Suillus*, Fig. 2d) and associated very low general root diversity (Fig. 1g-h) is consistent also with pre-HTP sequencing era studies of undisturbed *P. sylvestris* roots (Jonsson *et al*., 1999).

It is possible that differences in disturbance is an explanation for the difference in *Mycena* infection levels, i.e. that *Mycena* root invasion should be seen as a largely opportunistic feature of plants that are young, disturbed or otherwise vulnerable. This is also consistent with what is generally known to facilitate attacks from known fungal parasites (Walters, 2011). We speculate that this disturbance effect could also apply within species: e.g. grazing by herbivores passing by at one location would be likely to impact multiple individuals close by each other. The clumped (uneven) distribution of *Mycena* infections levels in *B.vivipara* can thus be interpreted as offering some support for this theory.

An important issue (question 4) is what is the basis of interaction when *Mycena* invades a host root. It was not possible to determine i the *Mycena* carpophores sampled for analysis of stable isotopes were free-living or from mycelia associated with a plant, Although the average value of Mycena suggested a saprotrophic lifestyle, there were several individual collections with profiles that suggested an alternative mode of nutrition. Most clearly, two collections of *Mycena pura* in Finse have isotopic profiles that resemble that of mycorrhizal fungi., Interestingly, in the study by Thoen *et al*., (2020) the culture of *M. pura* which could transfer ^32^P to its plant host was grown from the *M. pura1* sample from Finse.

In addition, the isotopic profiles of *M. pura* and its close relatives *M. rosea2* and *M. diosma* of the section (Calodontes) from Gribskov (in Denmark) are not far from those of the ectomycorrhizal *Russula* or *Cortinarius*.(Fig. 4 However, the *M. pura* from Svalbard and Vettakollen (in mainland Norway) and a *M. rosea1* from Gribskov had isotopic signatures closer to the expected profiles for saprotrophs. The members of the *Mycena* section Calodontes are notably hard to grow in culture which has previously led to speculations about their nutrition (Perreau *et al*., 1992; Boisselier-Dubayle *et al*., 1996; Harder *et al*., 2010; Harder *et al*., 2012). However, it must be noted that *Mycena pura* was rarely found in the root samples and only constituted a significant (>10%) fraction of the root community in one single *B. vivipara* individual (Fig. S5). None of our collections of *M. galopus* displayed similar intraspecific variations and/or mycorrhizal-like patterns in their isotopic profiles as one might have expected based on Grelet *et al*. (2017) or Thoen *et al*. (2020). Whether this is a coincidence for this study or not must be left for further research to explore.

Overall, the variable isotopic patterns of *Mycena* from the field are broadly consistent with the interspecific and intraspecific variation observed in the interactions between *Mycena* and birch roots in the growth experiments by Thoen *et al*. (2020), where different species and conspecific strains displayed harmful, neutral/endophytic, or beneficial interaction phenotypes *in vitro*. In the light of the emerging discoveries of variable ecologies among several fungal genera, these findings highlight the need for more targeted organism-level research on multiple individuals of fungal species to obtain a more comprehensive picture of the possible ecological versatilities. If fungal ecology is versatile not only below the genus, but also below the species level, then this may lead to reconsideration of the high importance ascribed to nutrition as a decisive taxon-delimiting trait (as in *Serpulaceae* (Skrede *et al*., 2011) or *Clavariaceae* (Birkebak *et al*., 2013)). Redhead *et al*. (2016) proposed to split the monophyletic *Amanita* sensu lato into ectomycorrhizal *Amanita* sensu stricto and a new saprotrophic *Saproamanita*, precisely to make ecological annotation in molecular studies easier, but this would be unnecessary with greater appreciation of ecological versatility. This has important implications for the widely applied approach in HTP-sequencing/metabarcoding plant root studies where annotating ecology to a sequence with genus-level based taxonomy could be misguided.

A question was raised by Vohník (2020) concerning the high numbers of *Mycena/Clavaria* sequences recovered from *C. tetragona* by Lorberau *et al*. (2017) suggesting that they might be explained by a lack of root cleaning/washing. While this is an issue that should not be ignored, all root samples included in the present study were either serially washed and/or surface sterilised as standard, including (Lorberau *et al*. (2017). It is therefore very unlikely that the recovery of large numbers of *Mycena* reads (and other saprotrophs) is purely due to mycelia living commensally on the root surface.

Until now, most data on the occurrence of saprotrophic fungi inside plant roots has arisen as a by-product of other research, and the sampling for these metabarcoding datasets here were not originally designed to investigate this question or to be analysed together. The differences (Fig 1) in average *Mycena* infection levels between 9 of the 10 host plants in our sample should be interpreted with the caution warranted by differences in sample sizes and site variations, and in comparing two ITS regions (Harder *et al*., 2013). The reverse primer (ITS2_r) of the ITS1 primer set has a terminal mismatch with 99% of all *Mycena* species (Tedersoo and Lindahl, 2016), which suggests that *Mycena* content in the ITS1 dataset could be underestimated. If true, this would merely strengthen our overall conclusions about *Mycena* as an overlooked but significant root invading genus; however, more studies directly targeting supposedly saprotrophic or endophytic (non-mycorrhizal) fungi in roots are certainly desirable.

To test our hypothesis that *Mycena* (or other saprotrophic) root infections are a result of opportunistic invasion under disturbance-related circumstances, future targeted metabarcoding root studies should directly analyse roots of multiple host species of different age and disturbance levels in the field in order to identify particular factors that may affect root invasion. Annotation databases should be continuously updated to reflect our best taxonomic knowledge<u>, and</u> further efforts should be undertaken to identify fungal OTUs or ASVs beyond the genus level, which may require more attention to detail than relying on SINTAX/RDP classification based on even the best possible databases.

Most importantly, more studies on direct ecological interactions between particular hosts and known fungal species are needed; both resynthesis host-fungus experiments with studies of nutrient and C transfer between the symbionts, and stable isotope studies in the field that specifically target saprotrophic taxa.

## Conclusions

The investigation of the trophic status of genus Mycena using sequence data from wild plant roots and 15N and 13C stable isotope signatures yielded the following: **1)** In nine of ten analysed herbaceous and ericaceous plants and tree mycorrhizal host plants from temperate, alpine and arctic environments, *Mycena* was consistently present in living plant roots across species and in different environments, but other saprotrophic taxa were only occasionally present; **2)** *Mycena* infections were not generally more prevalent in Arctic environments or at higher altitudes, but we hypothesise that infection may be more prevalent under conditions of disturbance; **3)** The ability to invade living plant roots is a feature of multiple *Mycena* species that do not discriminate between plant hosts, and **4)** The stable isotopic data on carpophores suggested that, although the genus *Mycena* is indeed mostly saprotrophic, strains of certain *Mycena* species can display an ecological versatility in the field and exchange nutrients with plants, consistent with previous results from *in vitro* resynthesis experiments.

The evidence that fungal trophic modes may be variable on the species level, and that within a large genus such as *Mycena* there may be several potential trophic options in addition to pure free-living saprotrophy, raises intriguing questions about the general understanding and study of fungal ecology. More research directly targeting root-associated fungi with unclear or unknown ecologies is required to resolve these questions. This study highlights the importance of continued detailed studies on interactions among organisms at the species level in order enhance data usage from broad, environmental metabarcoding approaches to community characterisation.

## Experimental procedures

### Sample site and sample descriptions

*Betula pubescens* roots were collected at the RSPB Nature Reserve at Corrimony in north-west Scotland in August 2008. The trees were regenerating saplings at a maximum 1 m in height, growing on moorland within heather-dominated vegetation on a site previously browsed by sheep and deer. Roots samples (supporting 100-200 ECM tips) were taken from the trees by direct tracing fine roots from the main laterals. Roots from 5 trees from within a block were pooled to give one single sample.

Roots from *Salix herbacea, Betula nana, Arctostaphylos. alpina*, and 8 *A. uva-ursi* from an original biogeography study were collected from mountains across Scotland (Fig S1) (Hesling and Taylor 2013).

The remaining 68 sample of *A. uva-ursi* roots were from an altitudinal gradient study (and 9 additional *P.sylvestris* samples in addition to those from Jarvis *et al*. (2015) in this study), collected June-July 2011 in the Invereshie-Inshriach National Nature Reserve in the north-west of the Cairngorms National Park in Scotland (Figs. S1, S2). Samples came from 9 elevation transects from 450-850 masl on a *Calluna*-*Arctostaphylos* subalpine heath with scattered Scots pine trees up until the tree limit at ~650 masl. This was in close proximity to the *P. sylvestris* forest studied in Jarvis *et al*. (2015)

The previously published datasets of *B. vivipara, S.polaris, D. octopetala* and *C.tetragona* were all collected in Arctic and Alpine tundra above the treeline in Arctic Norway, Iceland and Austria, and from grassland below the treeline in Scotland. For more details on the previously published data, we refer to the original publications. A more detailed description of the plant species targeted and the sample sites for the new data can be found in the supplementary data.

### Preparation of roots and old amplicon libraries for previously published ITS2 datasets

For the three new ITS2 datasets/454 runs representing 5 of 7 host species (see bioinformatics below), roots were sampled and cleaned under a dissection microscope to remove visible soil debris, woody and non-target species’ roots, then lyophilized in 2 ml tubes and milled using a steel bead in a mixer mill (RETSCH, Düsseldorf, Germany). Dry weight for DNA extraction was adjusted for each sample so that extracted mass was proportional to total sample dry weight: *extract weight* (mg) = (42.50 × *total sample dry weight* (mg)) + 47.98. DNA was extracted using 96 well, DNeasy Plant Minikits (QIAGEN, Hilden, Germany).

PCR amplification of the ITS2 region was conducted on a 2720 Thermal Cycler (Life Technologies, Carlsbad, CA, USA) in 10 μl reactions: 5 μl diluted template; 40 μM of each nucleotide; 0.55 mM MgCl_2_; 40 nM ITS7A primer (Ihrmark *et al*. 2012); 40 nM ITS 4 primer with a 3’ 8 bp tag (unique by ≥2 bp between samples); and 0.005 U/μl polymerase (DreamTaq Green, Thermo Scientific, Waltham, MA, USA) in buffer. Cycling parameters were: 94 °C for 5 min then 25, 30 or 35 cycles at 94 °C for 30 s; 57 °C for 30 s; 72 °C for 30 s; with a final extension of 72 °C for 10 min. PCR products were checked using gel electrophoresis (dilutions/cycles adjusted if products were out with the range 1-10 ng μl^-1^), then purified using AMPure 96 (Beckman Coulter, Brea, USA). DNA concentrations were established using a Qubit 2.0 fluorometer (Invitrogen, Paisley, UK), samples combined in equal molar proportion, further purified using GeneJET PCR Purification (Thermo Scientific, Waltham, USA) and lyophilized. Adaptor ligation, 454-sequencing and sequence adapter trimming were performed by the NERC genomics facility (Liverpool, UK) on one pico-titre plate using the GL FLX Titanium system (Roche, Basel, Switzerland).

## *Mycena* database

For identification of *Mycena* sequence data to species level, we first generated 151 new sequences from herbarium specimens and personal collections, using the ITS1F/ITS4 primers and the PCR protocol of Gardes and Bruns (1993). All *Mycena* ITS full-length sequences from GenBank and from the UNITE database (1099 sq) were extracted. Sequences not identified to species level, which did not cover the regions amplified by the ITS1F-2/ITS3-4 primer target regions, and which were not inside the *Mycena* sensu stricto clade (Fig. 3), or duplicates between both databases were discarded. Additionally, 14 complete *Mycena* sequences in GenBank were also discarded, as these were deemed to be misidentified (see table S15), most of those from (Hofstetter *et al*. 2019). The final database comprised 576 high-quality sequences with 136 named *Mycena* species, 89 of which with ≥2 sequences.

Sequences were aligned with the FFT-NS-i algoritm in MAFFT v 5 (Katoh and Standley, 2013). The complete (628 bp) and annotated *M. pura* EU517504 sequence was used to identify the ITS1 and ITS2 regions in the alignment.

## Bioinformatics

From the previously published studies of *B. vivipara*, a high-quality dataset of 119 054 sequences was compiled from Blaalid et al (2012), 191,099 from Yao *et al*. (2013),157 181 from Botnen *et al*. (2014), 244 523 from Blaalid *et al*. (2014) 272 595 sequences from Mundra *et al*., (2015) 249 888 from Davey *et al*. (2015), and 132912 from Botnen *et al*. (2019), making a total of 1095997 ITS1 sequences for clustering into OTUs/ASVs.

For the ITS2 dataset, we first analysed the two previously published studies of Jarvis et al (2015) and Lorberau et al (2017), and obtained 175829 and 1952314 sequences, For the three unpublished ITS2 454 runs, 327480 raw reads were obtained on a run with 104 *Betula pubescens* samples; 494187 raw reads on a run combining altitude and biogeography samples, and 232125 for a run with 16 1st year biogeography samples. After denoising, chimera check, length, primer/base pair match and quality controls, 121587, 326380 and 154121 high-quality reads remained, respectively. In total, this amounted to 2730231 high-quality ITS2 sequences of all fungal ecological groups. These were then used for clustering into OTUs/ASVs.

The OTUs/ASVs were classified taxonomically with the non-Bayesian SINTAX classifier (Edgar, 2016) using the 8.2 utax eukaryote reference database (Abarenkov *et al*., 2020).

QIIME (Caporaso *et al*., 2010) 1.9.1 pipeline was employed for the three unpublished 454 runs through the same steps as in Jarvis *et al*., (2015) until the OTU clustering step. We retained those with a sequence length 200-550 bp, only 100% match to in primer/tag sequences, passed chimera check in UCHIME(Edgar *et al*., 2011), a sliding window quality check of 50 bp applied to identify low-quality regions (average Phred score < 25). The resulting fasta files from the individual ITS1 and ITS2 runs were combined, and clustered first into OTUs at 97% similarity using vsearch (Rognes *et al*., 2016) and its usearch_global command function, and then into ASVs using the standard settings in UNOISE (Edgar, 2016). The R decontam package (Davis *et al*., 2018) with the default settings to remove likely contaminants based on the negative controls on a per sample basis for each of the 6 different datasets in the ITS1 part, and on the single negative control samples in the *Cassiope tetragona* ITS2 dataset (no negative controls were sequenced in Jarvis *et al*. (2015) nor in any of the new ITS2 datasets). Non-fungal sequences (P<0.95), and OTUs and ASVs with respectively <10/<8 counts were removed. Finally, sampling saturation was assessed with the iNEXT package (Hsieh *et al*., 2016), (see also Fig. S5), and all samples not meeting a coverage-based completeness (Chao and Jost, 2012) of 97% were discarded.

### Collection of samples for isotope analysis

Fungal carpophores, host plants and soil were collected from 5 different regions: Svalbard (arctic Norway); Finse/Hardangervidda (alpine central Norway), Vettakollåsen (boreal forest) in southeastern Norway; Solhomfjell National Park (boreal forest) in South Norway, and Gribsø /Gribskov (North Zealand, Denmark), in a beech-dominated broadleaf forest patch (Fig. S1) in 2015 and 2016. For more information and geographic coordinates of the field locations, see Fig. S1 and legend. In Svalbard, the collection sites spanned several similar valleys on the southern banks of Isfjorden, with the sites separated by up to ~60 km (Fig. S1); for the other four remaining collection sites, samples were taken from an area that extended over no more than 1 km^2^.

Fungal carpophores, soil samples and plants were dried with continuous airflow for 12-36 hours at 70 °C until dry. Plants and fungi were identified morphologically, and *Mycena* furthermore by ITS sequences. For Svalbard, some additional fungal samples were taken from dried mushroom specimens kept at the herbarium at Tøyen at the Natural History Museum in Oslo. It was assumed that individual carpophores collected within a distance of <50 centimeters between them originated from the same mycelium. Conspecific *Mycena* carpophores sampled from larger distances were treated as separate samples. Whenever possible, collections from a given site were triplicated or at least duplicated, using separate fruit bodies from the same collection. Fungi were divided into the three categories “ectomycorrhizal”, “saprotrophic” or *“Mycena”*. For every 36 samples analysed, internal replicates of material from two samples from the same fruitbody was used to verify consistent machine functioning. Soil samples were taken from top-soil (A horizon, 0 cm) and from mineral soils in ~50 cm depths. We sampled soil from three different locations on the different sites. Plant samples were all taken from leaves.

### Stable isotope analysis

Dried samples (plants, fungi, soil) were ground by hand, weighed (see supplementary table TS4) into 5 x 9 mm tin capsules (Sercon), closed and compressed. Samples consisted of 5 mg of fungi/plant, 10 mg of topsoil, or 20 mg of 50 cm depth soil. Samples were analysed for δ^13^C, δ^15^N, % C, and % N by continuous flow with a Costech ECS4010 elemental analyser (Costech Analytical Technologies Inc, Valencia, California) coupled with a DELTAplus XP isotope ratio mass spectrometer (Thermo Scientific, Bremen, Germany) at the University of New Hampshire Stable Isotope Laboratory. All carbon and nitrogen isotope data are reported in delta notation according to this equation: δX = [(R_sample_/R_standard_) - 1] × 1000 where X is ^13^C or ^15^N and R is the ratio ^13^C/^12^C or ^15^N/^14^N. All δ^13^C and δ^15^N values were normalised on VPDB (δ^13^C) and AIR (δ^15^N) reference scales with the following internationally calibrated standards and values: IAEA CH6 (210.45%), CH7 (232.15%), N1 (0.4%) and N2 (20.3%). Laboratory working standards included NIST 1515 (apple leaves), NIST 1575a (pine needles) and tuna muscle, as well as a *Boletus* quality control.

#### Statistics/graphics

Stepwise multiple regression models of fungal δ^15^N and δ^13^C were analyzed with genus, site, and %N as the independent variables. Because of the declining δ^13^C of atmospheric carbon dioxide, year was also included as a continuous factor in the δ^13^C regression. Genus and site were categorical variables and year and %N were continuous variables. These statistical analyses were carried out in JMP 13 Pro (SAS Institute, Middleton, Massachusetts, USA). Models that minimized the Bayesian Information Criterion (BIC) were selected. The variance inflation factor (VIF) of each model factor was also calculated, which measures multicollinearity. This approach allowed a test of whether *Mycena* generally grouped with saprotrophic or ectomycorrhizal genera without *a priori* setting up specific contrasts among *Mycena*, saprotrophic genera, and ectomycorrhizal genera.

All other statistics were done in R using’phyloseq’ 1.19.1 R package (McMurdie and Holmes, 2013) for combining and rearranging OTU tables and taxonomy information, and the heatmap.2 function from the ‘gplots’ package (Warnes *et al*., 2016) for visualising heatmaps. We applied a sequential ANOVA for Fig. 1a-d at the 0.05 significance threshold with the Scheffe post-hoc test correction for multiple comparisons, using the ‘agricolae’ package (de Mendiburu, 2020).

## Phylogenetics

The ITS phylogeny (Fig. 3) was constructed by first aligning a selected high-quality subset of 89 ITS full-length sequences with the Q-ins-i algoritm in MAFFT(Katoh and Standley, 2013) for a final alignment of 1502 positions (gaps included), and then running a maximum likelihood with 1000 bootstrap replications in RaxML (Stamatakis, 2014), saving branch lengths. Then, the 20 + 21 *Mycena* ITS1 and ITS2 OTUs were added the, the Q-ins-i alignment redone, and the OTUs mapped to the branches using the EPA algoritm (Barbera, 2019). The tree was visualised in FIGTREE v. 1.4.3 (http://tree.bio.ed.ac.uk/software/figtree).

### Sequences and data

MiSeq/454 files are found at the respective sources listed in Table S16. Sanger sequences can be accessed through GenBank/UNITE, see Table S17 for accession numbers. R scripts and downstream analysis files can be obtained from C.B. Harder upon request.

## Acknowledgements

We thank Karl-Henrik Larsson and Arne Aronsen for provisions of specimens from the Natural History Museum of Oslo, and for help with identification of field specimens from Svalbard. We further thank Cecilie Mathiesen and Mikayla Jacobs for technical assistance in the laboratory, to Brendan J. Furneaux for a valuable input to the R script, and to the curators of H, TUR and OULU. The Mycena ITS sequences originating from the specimens deposited in H, TUR and OULU were produced as part of the Finnish Barcode of Life Project (FinBOL) funded by the Ministry of Environment, Finland (YM23/5512/2013), Otto A Malm’s Donationsfond, and the Kone Foundation.

We thank the European Commission (grant no. 658849) and the Carlsberg Foundation (grant no. CF18-0809) for grants to C.B. Harder that made this research possible.

## Conflicts of interest

The authors declare no conflicts of interest.

## Notes

### Competing Interest Statement

The authors have declared no competing interest.

