## Supplementary data for "*Mycena* species can be opportunist-generalist plant root invaders"

| | ITS1 | | ITS2 | |
|---|---|---|---|---|
| | Seqs | OTUs | Seqs | OTUs |
| Fungi (0.95) | 992852 | 1911 | 1173834 | 1691 |
| Raw in OTU table (n≥10) | 989946 | 1302 | 1170751 | 1057 |
| After decontam | 915747 | 1206 | 1016235 | 1044 |
| Final(97% coverage) | 889290 | 1193 | 992890 | 1032 |

Table S1. Total sequence count + OTUs for ITS1 + ITS2 datasets at each OTU table trimming step.[1]Non-target samples discarded.

| | ITS1 | | ITS2 | |
|---|---|---|---|---|
| | Seqs | zOTUs | Seqs | zOTUs |
| All zotus[1] | 1061048 | 3135 | 2689730 | 1880 |
| Fungi (0.95) | 1024007 | 3094 | 1210397 | 1705 |
| After decontam | 997077 | 3012 | 1107636 | 1685 |
| Final(97% coverage) | 967316 | 2772 | 1083789 | 1559 |

Table S2. Total sequence count + zotus (n=8) for ITS1 + ITS2 datasets at each zotu table trimming step. [1]Non-target samples and "zotus"/ASVs with <8 copies discarded.

| | Bistorta vivipara (n=519)[1] | Dryas octopetala (n=22) | Salix polaris (n=20)[2] |
|---|---|---|---|
| Fungi (0.95) | 917667 | 41314 | 33871 |
| Raw in OTU table (n≥10) | 914888 | 41272 | 33786 |
| After decontam | 843137 | 39880 | 32730 |
| Final (97% coverage) | 803649 | 39880 | 30069 |
| Sequences per sample | min=251, avg=1523.6, max=7680 | min=382, avg=1812.7, max=3735 | min=449, avg=1551.2, max=3516 |

Table S3. ITS1 data quality filtering steps (OTUs). For *Bistorta vivipara*, 80 samples of the original 599 that had too few sequences to meet the coverage threshold of 97% in both the OTU and zotu datasets were discarded. [2]1 *Salix polaris* sample was discarded for the same reason.

| | Bistorta vivipara (n=519)[1] | Dryas octopetala (n=22) | Salix polaris (n=20)[2] |
|---|---|---|---|
| Fungi (0.95) | 984812 | 44702 | 36145 |
| After decontam | 919672 | 43286 | 34119 |
| Final (97% coverage) | 908621 | 43286 | 33141 |
| Sequences per | min=251, avg=1519.4, | min=382, avg=1812.7, | min=449, avg=1546.7, |

| | | | |
|---|---|---|---|
| sample | max=7680 | max=3730 | max=3516 |

Table S4. ITS1 data quality filtering steps (zotus). [1]For *Bistorta vivipara*, 80 samples of the original 599 that had too few sequences to meet the coverage threshold of 97% in both the OTU and zotu datasets were discarded. [2]1 *Salix polaris* sample was discarded for the same reason.

| Plant host | Arctostaphylos alpine (n=10) | A.uva-ursi (n=76) [1] | B.nana (n=8) | Betula pendula (n=81)[2] | Pinus sylvestris (n=41) | Salix herbacea (n=7) | Cassiope tetragona (n=15) |
|---|---|---|---|---|---|---|---|
| Fungi (0.95) | 60168 | 249638 | 50124 | 107463 | 212727 | 25604 | 468110 |
| Raw in OTU table (n≥10) | 59878 | 248708 | 49945 | 106620 | 212589 | 25322 | 467689 |
| After decontam | 59829 | 248428 | 49927 | 106549 | 212577 | 25298 | 313627 |
| Final (97% coverage) | 59829 | 241418 | 49927 | 90214 | 212577 | 25298 | 313627 |
| Sequences per sample | min=2173 avg=5998.2 max=9437 | min=665, avg=3047.8 max=10187 | min=3157 avg=6243.1max=10646 | min=536 avg=1019.2 max=2240 | min=669 avg=5185.7 max=19286 | min=1996 avg=3627.6 max=7025 | min= 420 avg=20911.8 max=56456 |

Table S5. ITS2 data quality filtering steps (OTUs). For *Arctostaphylos uva-ursi* and *Betula pendula*, respectively 6 and 23 of the original 82 and 104 samples had too few sequences to meet the coverage threshold of 97% in both the OTU and zotu datasets and were discarded.

| Plant host | Arctostaphylos alpine (n=10) | A.uva-ursi (n=76) [1] | B.nana (n=8) | Betula pendula (n=81)[2] | Pinus sylvestris (n=41) | Salix herbacea (n=7) | Cassiope tetragona (n=15) |
|---|---|---|---|---|---|---|---|
| Fungi (0.95) | 61718 | 254512 | 51229 | 112588 | 219210 | 25535 | 485605 |
| After decontam | 61703 | 254557 | 51182 | 109810 | 219124 | 25497 | 385763 |
| Final (97% coverag | 61703 | 247091 | 51182 | 93429 | 219124 | 25497 | 385763 |

| e) | | | | | | | |
|---|---|---|---|---|---|---|---|
| Sequences per sample | min=2157 avg=6170.3 max=9842 | min=846, avg=3272 max=10298 | min=3151 avg=6397.8 max=10793 | min=550 avg=1153.4 max=2388 | min= 670 avg=5344.5 max=20526 | min= 2022 avg=3642.4 max=7083 | min= 563 avg=25717.5 max=61829 |

Table S6. ITS2 data quality filtering steps (zotus). For *Arctostaphylos uva-ursi*, and *Betula pendula*, respectively 6 and 23 of the original 82 and 104 samples had too few sequences to meet the coverage threshold of 97% in both the OTU and zotu datasets and were discarded.

| Host plant | $R^2$ | P-value |
|---|---|---|
| Arctostaphylos alpine (n=10) | 0.2433 | 0.08393 |
| A.uva-ursi (n=76) [1] | 0.02791 | 0.07193 |
| B.nana (n=8) | 0.2525 | 0.1163 |
| Betula pendula (n=81)[2] | -0.00519 | 0.4861 |
| Pinus sylvestris (n=41) | 0.1176 | 0.0161 |
| Salix herbacea (n=7) | 0.05233 | 0.3007 |
| Cassiope tetragona (n=15) | -0.06255 | 0.6818 |
| Bistorta vivipara (n=519)[1] | 0.001058 | 0.2139 |
| Dryas octopetala (n=22) | -0.04172 | 0.6944 |
| Salix polaris (n=20) [2] | -0.01962 | 0.4361 |

Table S7. *Mycena* infection load (% reads of total) correlations with sampling depth.

| **ITS1 OTUs** | ***Mycena* species identification** | **Host plant** | **ITS1 ASVs** | ***Mycena* species identification** | **Host pla** |
|---|---|---|---|---|---|
| OTU_614 | *M.sp_monticola((0.915)* | Bv | ASV_1232 | *M.sp_monticola((0.915)* | Bv |
| OTU_201 | *M.sp.pasvikensis(0.95)* | Bv | ASV_298 | *M.sp_pasvikensis(0.95a)* | Bv |
| - | - | | ASV_1358 | *M.sp_pasvikensis(0.95b)* | Bv |
| - | - | | ASV_3028 | *M.sp_plumbea(0.916)* | None (>1%) |
| OTU_505 | *M.sp.rebaudengoi(0.93)* | Bv | ASV_962 | *M.sp_rebaudengoi(0.93)* | Bv |
| OTU_448 | *M.sp.rebaudengoi(0.95)* | Bv | ASV_844 | *M.sp_rebaudengoi(0.95a)* | Bv |
| - | - | | ASV_1298 | *M.sp_rebaudengoi(0.95b)* | Bv |
| - | - | | ASV_2179 | *M.sp_tenax(0.92)* | None |

| | | | | | |
|---|---|---|---|---|---|
| | | | | | (>1%) |
| OTU_124 | *M.alb/oli/citr* | Bv,Do | ASV_147 | *M.alb/oli/citr* | Bv,Do |
| OTU_1828 | *M.cinerella* | Bv | ASV_1179 | *M.cinerella1* | Bv |
| - | - | | ASV_2044 | *M.cinerella2* | Bv |
| - | - | | ASV_246 | *M.cinerella3* | Bv |
| OTU_274 | *M.concolor* | Bv | ASV_804 | *M.concolor1* | Bv |
| - | - | | ASV_1638 | *M.concolor2* | Bv |
| - | - | | ASV_1705 | *M.concolor3* | Bv |
| - | - | | ASV_1559 | *M.concolor4* | Bv |
| - | - | | ASV_1777 | *M.concolor5* | Bv |
| - | - | | ASV_2131 | *M.concolor6* | Bv |
| - | - | | ASV_1710 | *M.concolor7* | Bv |
| OTU_90 | *M.epipterygia1* | Bv | ASV_346 | *M.epipterygia1* | Bv |
| OTU_618 | *M.epipterygia2* | Do | ASV_1237 | *M.epipterygia2* | Bv |
| OTU_762 | *M.gal/meg* | Bv | ASV_1631 | *M.gal/meg* | Bv |
| OTU_631 | *M.galopus* | Bv | ASV_1274 | *M.galopus* | Bv |
| OTU_314 | *M.metata* | Bv | ASV_540 | *M.metata* | Bv |
| OTU_118 | *M.pasvikensis* | Bv | ASV_175 | *M.pasvikensis1* | Bv |
| - | - | | ASV_638 | *M.pasvikensis2* | Bv |
| OTU_426 | *M.pura* | Bv | ASV_799 | *M.pura* | Bv |
| OTU_1169 | *M.rebaudengoi* | None (>1%) | ASV_2835 | *M.rebaudengoi* | None (>1%) |
| OTU_1204 | *M.sanguinolenta* | Bv | ASV_2963 | *M.sanguinolenta* | Bv |
| OTU_543 | *M.leptocephala* | Bv | ASV_1053 | *M.leptocephala* | Bv |
| OTU_21 | *M.strob/aur* | Bv,Do,Sp | ASV_19 | *M.strob/aur1* | Bv,Do,S[illegible] |
| - | - | | ASV_440 | *M.strob/aur2* | Bv |
| - | - | | ASV_543 | *M.strob/aur3* | Bv |
| - | - | | ASV_724 | *M.strob/aur4* | Bv |
| - | - | | ASV_2904 | *M.strob/aur5* | None (>1%) |
| OTU_51 | *M.stylobates1* | Bv,Do | ASV_50 | *M.stylobates* | Bv,Do,S[illegible] |
| OTU_598 | *M.stylobates2* | None (>1%) | ASV_1188 | *M.stylobates* | Bv |
| NA | (No corresponding Mycena-OTU) | | ASV_1838 | *M.sp_cinerella(0.93)* | None (>1%) |
| NA | (No corresponding Mycena-OTU) | | ASV_1755 | *M.sp_cinerella(0.95)* | None (>1%) |
| NA | (No corresponding Mycena-OTU) | | ASV_1600 | *M.sp_cinerella(0.96)* | None (>1%) |
| NA | (No corresponding Mycena-OTU) | | ASV_2791 | *M.sp_citrinomarginata(0.95)* | None (>1%) |
| NA | (No corresponding Mycena-OTU) | | ASV_2659 | *M.sp_epipterygia(0.93)* | None (>1%) |
| NA | (No corresponding Mycena-OTU) | | ASV_2167 | *M.sp_metata(0.91)* | None |

| | | | | |
|---|---|---|---|---|
| | | | | (>1%) |
| NA | (No corresponding Mycena-OTU) | ASV_1895 | *M.sp_metata(0.93)* | None (>1%) |
| NA | (No corresponding Mycena-OTU) | ASV_950 | *M.sp_metata(0.97)* | Bv |

Table S8. Total list of OTUs and ASVs in the ITS1 data identified as *Mycena* by clustering into new OTUs at 3% similarity with the *Mycena* database. Thick dark lines separate delimitations of ASV groups that clustered together with a particular OTU from the ITS1 dataset. The bottom 8 ASVs did not cluster together with any OTUs. For sequences that did not cluster together with any named species in the *Mycena* database ("M.sp_[epithet](number)") and thus could not be identified to the species level, species epithet followed by number in parentheses denote sequence similarity to its most similar species in the database. For all OTUs/ASV, only plant hosts where at least 1% of the reads in at least one sample represents the particular OTU/ASV are listed under "Host plant" columns. "None (>1%)" then means that no hosts met this criterion.
Host plant abbreviations: Bv=*Bistorta vivipara*, Do=*Dryas octopetala*, Sp=*Salix polaris*.Mycena abbreviations: M.strob/aur= *M.strobiluroides/aurantiomarginata*, M.gal/meg=*M.galericulata/megaspora,* M.alb/oli/citr= *M.albidolilacea/olivaceomarginata/citrinomarginata*.

| **ITS2 OTUs** | ***Mycena* species identification** | **Host plant** | **ITS2 ASVs** | ***Mycena* species identification** | **Host plant** |
|---|---|---|---|---|---|
| OTU_232 | *M.aciculata* | Bp | ASV_348 | *M.aciculata* | Aa |
| OTU_73 | *M.alb/oli/citr* | Bp,Ct | ASV_115 | *M.alb/oli/citr1* | Ct |
| - | - | - | ASV_548 | *M.alb/oli/citr2* | None (>1%) |
| - | - | - | ASV_561 | *M.alb/oli/citr3* | Bp |
| - | - | - | ASV_1562 | *M.alb/oli/citr4* | None (>1%) |
| OTU_358 | *M.alexandri* | None (>1%) | ASV_698 | *M.alexandri1* | None (>1%) |
| - | - | - | ASV_770 | *M.alexandri2* | None (>1%) |
| OTU_194 | *M.amicta* | Bp | ASV_300 | *M.amicta1* | Bp |
| - | - | - | ASV_697 | *M.amicta2* | None (>1%) |
| OTU_20 | *M.cinerella* | Aa,Au,Bn,Bp,Sh | ASV_130 | *M.cinerella1* | Aa,Au,Sh |
| - | - | - | ASV_31 | *M.cinerella2* | Bp |
| OTU_4 | *M.epipterygia(OTU1)* | Aa,Au,Bn,Bp,Ct | ASV_7 | *M.epipterygia1* | Ct |
| - | - | - | ASV_9 | *M.epipterygia2* | Ct |
| - | - | - | ASV_63 | *M.epipterygia3* | Aa,Au,Bn,Bp |
| - | - | - | ASV_16 | *M.epipterygia4* | Aa,Au,Bn |

| | | | | | |
|---|---|---|---|---|---|
| | | | 9 | | |
| - | - | | ASV_179 | *M.epipterygia5* | Aa,Au |
| OTU_883 | *M.epipterygia(OTU2)* | None (>1%) | ASV_1577 | *M.epipterygia6* | None (>1%) |
| OTU_400 | *M.gal/meg* | None (>1%) | ASV_788 | *M.gal/meg* | Ct |
| OTU_10 | *M.galopus* | Aa,Au,Bn,Bp,Sh | ASV_114 | *M.galopus1* | Aa,Au,Bn |
| - | - | - | ASV_16 | *M.galopus2* | None (>1%) |
| - | - | - | ASV_25 | *M.galopus3* | Ct |
| OTU_185 | *M.latifolia* | Bp | ASV_1154 | *M.latifolia1* | None (>1%) |
| - | - | - | ASV_267 | *M.latifolia2* | Bp |
| OTU_6 | *M.leptocephala* | Aa,Au,Bn,Bp,Ct,Sh | ASV_15 | *M.leptocephala1* | Bp,Ct |
| - | - | - | ASV_113 | *M.leptocephala2* | None (>1%) |
| - | - | - | ASV_149 | *M.leptocephala3* | Bp |
| OTU_17 | *M.metata* | Aa,Au,Bn,Bp,Ct,Sh | ASV_58 | *M.metata1* | Ct |
| - | - | - | ASV_87 | *M.metata2* | Ct |
| - | - | - | ASV_95 | *M.metata3* | Aa,Au,Bn,Sh |
| - | - | - | ASV_177 | *M.metata4* | Aa,Au,Bn,Sh |
| - | - | - | ASV_195 | *M.metata5* | Bn,Ct |
| - | - | - | ASV_203 | *M.metata6* | Bp |
| OTU_33 | *M.pasvikensis* | Bp,Ct | ASV_85 | *M.pasvikensis1* | Ct |
| - | - | - | ASV_109 | *M.pasvikensis2* | Ct |
| - | - | - | ASV_783 | *M.pasvikensis3* | Bp |
| - | - | - | ASV_577 | *M.rebaudengoi* | Bp |
| OTU_948 | *M.rubromarginata* | None (>1%) | ASV_1492 | *M.rubromarginata* | None (>1%) |
| OTU_13 | *M.sanguinolenta* | Au,Bn,Bp,Ps,Sh | ASV_26 | *M.sanguinolenta1* | None (>1%) |
| - | - | - | ASV_49 | *M.sanguinolenta2* | Au |
| OTU_743 | *M.septentrionalis* | None (>1%) | ASV_1325 | *M.septentrionalis* | None (>1%) |

| | | | | | |
|---|---|---|---|---|---|
| OTU_536 | *M.capillaripes* | None (>1%) | ASV_511 | *M.capillaripes* | Au |
| OTU_127 | *M.strob/aur* | Bp,Ct | ASV_222 | *M.strob/aur1* | Ct |
| - | - | - | ASV_688 | *M.strob/aur2* | Bp |
| OTU_139 | *M.stylobates* | Bp,Ct | ASV_354 | *M.stylobates1* | None (>1%) |
| - | - | - | ASV_626 | *M.stylobates2* | None (>1%) |
| - | - | - | ASV_801 | *M.stylobates3* | Bp |
| - | - | - | ASV_927 | *M.stylobates4* | Ct |
| OTU_723 | *M.sp. flos-nivum(0.98)* | Bp | ASV_1246 | *M.sp. flos-nivum(0.98)* | Bp |
| OTU_1061 | *M.sp_subcana(0.97)* | None (>1%) | | (No corresponding Mycena-ASVs) | |
| NA | (No corresponding Mycena-OTU) | NA | ASV_186 | *M.sp1* | Au |
| NA | (No corresponding Mycena-OTU) | NA | ASV_223 | *M.sp2* | Bp |
| NA | (No corresponding Mycena-OTU) | NA | ASV_483 | *M.sp3* | Au |
| NA | (No corresponding Mycena-OTU) | NA | ASV_238 | *M.abramsii1* | Ct |
| NA | (No corresponding Mycena-OTU) | NA | ASV_502 | *M.abramsii2* | None (>1%) |
| NA | (No corresponding Mycena-OTU) | NA | ASV_1402 | *M.clavicularis1* | None (>1%) |
| NA | (No corresponding Mycena-OTU) | NA | ASV_187 | *M.clavicularis2* | Au |
| NA | (No corresponding Mycena-OTU) | NA | ASV_327 | *M.concolor* | None (>1%) |

Table S9. Total list of OTUs and ASVs in the ITS2 data identified as *Mycena* by clustering into new OTUs at 3% similarity with the *Mycena* database. Thick dark lines separate delimitations of ASV groups that clustered together with the particular OTU from the ITS2 dataset. The bottom 8 ASVs did not cluster together with any OTUs. For sequences that did not cluster together with any named species in the *Mycena* database ("M.sp_[epithet](number)") and thus could not be identified to the species level, species epithet followed by number in parentheses denote sequence similarity to its most similar species in the database. For all OTUs/ASV, only plant hosts where at least 1% of the reads in at least one sample represents the particular OTU/ASV are listed under "Host plant" columns. "None (>1%)" then means that no hosts met this criterion. Host plant abbreviations: Aa=*Arctostaphylos alpine*, Au=*A. uva-ursi*, Bn=*Betula nana*, Bp=*B.pendula*, Ct=*Cassiope tetragona*.Ps=*Pinus sylvestris*, Sh=*Salix herbacea*. Mycena abbreviations: M.gal/meg=*M.galericulata/megaspora,* M.alb/oli/citr= *M.albidolilacea/olivaceomarginata/citrinomarginata*.

| **Pairwise Comparison - *Mycena* infection level** | **difference** | **p-value** | **signif.** | **LCL** | **UCL** |
|---|---|---|---|---|---|
| *B.vivipara - D.octopetala* | -0.12511525 | 0.0000 | *** | -0.16518000 | -0.08505050 |
| *B.vivipara - S.polaris* | -0.14247594 | 0.0000 | *** | -0.18441846 | -0.10053342 |
| *D.octopetala - S.polaris* | -0.01736069 | 0.7533 | | -0.07422727 | 0.03950589 |
| **Pairwise Comparison -species richness at 97% cov** | **difference** | **p-value** | **signif.** | **LCL** | **UCL** |
| *B.vivipara - D.octopetala* | 0.1947128 | 0.9924 | | -3.697995 | 4.087421 |
| *B.vivipara - S.polaris* | -0.1377645 | 0.9965 | | -4.212917 | 3.937388 |
| *D.octopetala - S.polaris* | -0.3324773 | 0.9890 | | -5.857658 | 5.192703 |

Table S10. ITS1 data comparisons of *Mycena* infection load (top) and 97% coverage-based species richness (bottom) using Scheffes test for correction for multiple comparisons. Each component is a matrix with differences in observed means, LCL and UCL the lower/upper end point of the interval. Significant P-values (corrected for multiple comparisons) in bold, with asterisks *,** and *** denoting 0.05,0.01 and 0.001 levels.

| **Pairwise Comparison - *Mycena* infection level** | **difference** | **p-value** | **signif.** | **LCL** | **UCL** |
|---|---|---|---|---|---|
| *A.alpine - A.uva-ursi* | -0.0087233685 | 1.0000 | | -0.14649037 | 0.12904364 |
| *A.alpine - B.nana* | -0.1280569193 | 0.4426 | | -0.32232169 | 0.06620785 |
| *A.alpine - B.pendula* | -0.1760051850 | **0.0033** | ** | -0.31327687 | -0.03873350 |
| *A.alpine - C.tetragona* | -0.2300524341 | **0.0011** | ** | -0.39724893 | -0.06285594 |
| *A.alpine - P.sylvestris* | 0.0916919285 | 0.4902 | | -0.05275089 | 0.23613474 |
| *A.alpine - S.herbacea* | -0.0006946679 | 1.0000 | | -0.20252119 | 0.20113185 |
| *A.uva-ursi - B.nana* | -0.1193335508 | 0.2334 | | -0.27156018 | 0.03289308 |
| *A.uva-ursi - B.pendula* | -0.1672818165 | **0.0000** | *** | -0.23268567 | -0.10187797 |
| *A.uva-ursi - C.tetragona* | -0.2213290656 | **0.0000** | *** | -0.33703911 | -0.10561902 |
| *A.uva-ursi - P.sylvestris* | 0.1004152970 | **0.0039** | ** | 0.02105615 | 0.17977444 |

| | | | | | |
|---|---|---|---|---|---|
| *A.uva-ursi - S.herbacea* | 0.0080287006 | 1.0000 | | -0.15373684 | 0.16979424 |
| *B.nana - B.pendula* | -0.0479482656 | 0.9657 | | -0.19972678 | 0.10383024 |
| *B.nana - C.tetragona* | -0.1019955148 | 0.6220 | | -0.28129378 | 0.07730275 |
| *B.nana - P.sylvestris* | 0.2197488478 | **0.0010** | *** | 0.06145505 | 0.37804265 |
| *B.nana - S.herbacea* | 0.1273622515 | 0.5580 | | -0.08459799 | 0.33932249 |
| *B.pendula - C.tetragona* | -0.0540472491 | 0.8033 | | -0.16916711 | 0.06107261 |
| *B.pendula - P.sylvestris* | 0.2676971135 | **0.0000** | *** | 0.18920098 | 0.34619325 |
| *B.pendula - S.herbacea* | 0.1753105171 | **0.0235** | * | 0.01396660 | 0.33665443 |
| *C.tetragona - P.sylvestris* | 0.3217443626 | **0.0000** | *** | 0.19816122 | 0.44532751 |
| *C.tetragona - S.herbacea* | 0.2293577662 | **0.0061** | ** | 0.04189308 | 0.41682246 |
| *P.sylvestris - S.herbacea* | -0.0923865964 | 0.6560 | | -0.25987412 | 0.07510093 |
| **Pairwise Comparison -species richness at 97% cov** | **difference** | **p-value** | **signif.** | **LCL** | **UCL** |
| A.alpine - A.uva-ursi | 21.5162684 | 0.108 | | 3.094243 3 | 9.938294 |
| A.alpine - B.nana | 9.2362750 | 0.9396 | | -16.740558 3 | 5.213108 |
| A.alpine - B.pendula | 5.7823198 | 0.9662 | | -12.573473 2 | 4.138112 |
| A.alpine - C.tetragona | 46.7560333 | 0.0 | *** | 24.398735 6 | 9.113332 |
| A.alpine - P.sylvestris | 58.5593634 | 0.0 | *** | 39.244657 7 | 7.874070 |
| A.alpine - S.herbacea | 6.7373286 | 0.9897 | | -20.250652 3 | 3.725309 |
| A.uva-ursi - B.nana | -12.2799934 | 0.5532 | | -32.635542 | 8.075555 |
| A.uva-ursi - B.pendula | -15.7339487 | 0.0 | *** | -24.479667 - | 6.988231 |
| A.uva-ursi - C.tetragona | 25.2397649 | 0.0 | *** | 9.767168 4 | 0.712362 |
| A.uva-ursi - P.sylvestris | 37.0430950 | 0.0 | *** | 26.431293 4 | 7.654897 |
| A.uva-ursi - S.herbacea | -14.7789398 | 0.3972 | | -36.410019 | 6.852139 |
| B.nana - B.pendula | -3.4539552 | 0.9988 | | -23.749581 1 | 6.841671 |
| B.nana - C.tetragona | 37.5197583 | 0.001 | *** | 13.544227 6 | 1.495290 |
| B.nana - P.sylvestris | 49.3230884 | 0.0 | *** | 28.156246 7 | 0.489930 |
| B.nana - S.herbacea | -2.4989464 1. | 1.0 | | -30.841995 2 | 5.844102 |
| B.pendula - C.tetragona | 40.9737136 | 0.0 | *** | 25.580035 5 | 6.367393 |
| B.pendula - P.sylvestris | 52.7770437 | 0.0 | *** | 42.280642 6 | 3.273445 |
| B.pendula - S.herbacea | 0.9550088 | 1.0 | | -20.619691 2 | 2.529708 |
| C.tetragona - P.sylvestris | 11.8033301 | 0.3418 | | -4.722048 2 | 8.328708 |
| C.tetragona - S.herbacea | -40.0187048 | 0.0 | *** | -65.086240 - 1 | 4.951169 |
| P.sylvestris - S.herbacea | -51.8220348 | 0.0 | *** | -74.218249 - 2 | 9.425820 |

Table S11. ITS2 data comparisons of *Mycena* infection load (top) and 97% coverage-based species richness (bottom) using Scheffes test for correction for multiple comparisons. Each component is a matrix with differences in observed means, LCL and UCL the lower/upper end point of the interval. Significant P-values (corrected for multiple comparisons) in bold, with asterisks *,** and *** denoting 0.05,0.01 and 0.001 levels.

| **Environmental parameter** | **$R^2$** | **P-value** |
|---|---|---|
| Precipitation Seasonality | -0.003311 | 0.953 |
| Mean Diurnal Range | -0.003268 | 0.8988 |
| Min_Temperature_of_Coldest_Month | -0.002958 | 0.7411 |
| Max_Temperature_of_Warmest_Month | -0.002869 | 0.7124 |
| Precipitation_of_Driest_Quarter | -0.002614 | 0.645 |
| Precipitation_of_Wettest_Quarter | -0.001026 | 0.4067 |
| Temperature_Annual_Range | -0.002818 | 0.6974 |
| Precipitation_of_Wettest_Month | -0.00216 | 0.555 |
| Mean_Temperature_of_Coldest_Quarter | -0.003223 | 0.8633 |
| Mean_Temperature_of_Warmest_Quarter | -0.003283 | 0.9137 |
| Annual mean preciitation | -0.001208 | 0.426 |
| Annual mean temperatue | -0.002059 | 0.5383 |
| Latitude | -0.002843 | 0.7911 |
| Longitude | 0.0007986 | 0.2621 |

Table S12. Environmental correlation tests for B. vivipara with *Mycena* infection levels (% reads of total).

| **Response d15N, BIC min., genus-place-%N** | | | | | |
|---|---|---|---|---|---|
| RSquare | 0,571 | | | | |
| RSquare Adj | 0,561 | | | | |
| Root Mean Square Error | 2,978 | | | | |
| Mean of Response | 0,698 | | | | |
| Observations (or Sum Wgts) | 253 | | | | |
| Analysis of Variance | | | | | |
| Source | DF | Sum of Squares | Mean Square | F Ratio | |
| Model | 6 | 2903,09 | 483,85 | 54,56 | |
| Error | 246 | 2181,44 | 8,87 | | Prob > F |
| C. Total | 252 | 5084,53 | | <.0001 | |
| Parameter Estimates | | | | | |
| Term | %Variance | Estimate | Std Error | Prob >\|t\| | VIF |
| Intercept | -- | 1,53 | 0,26 | <.0001 | -- |
| Place{Gribskov&Solhomfjell-Finse&Vettakollen&Svalbard} | 8,4 | -1,26 | 0,19 | <.0001 | 1,03 |

| | | | | | |
|---|---|---|---|---|---|
| Type_genus{Trametes&Microcephale&Pluteus&Naucoria&Hygrophoropsis&Cystoderma&Lycoperdon&Rhodocollybia&Calvatia&Mycena&Galerina&Amanita&Helvella&Hypholoma-Lepista&Lactarius&Laccaria&Collybia&Ramaria&Russula&Inocybe&Cortinarius&Suillus&Rhizopogon&Hydnellum&Agaricus&Tricholoma&Hydnum} | 41,1 | -3,94 | 0,27 | <.0001 | 1,90 |
| Type_genus{Trametes&Microcephale&Pluteus&Naucoria&Hygrophoropsis&Cystoderma-Lycoperdon&Rhodocollybia&Calvatia&Mycena&Galerina&Amanita&Helvella&Hypholoma} | 3,8 | -1,71 | 0,38 | <.0001 | 1,65 |
| Type_genus{Lycoperdon&Rhodocollybia&Calvatia&Mycena-Galerina&Amanita&Helvella&Hypholoma} | 1,7 | -0,82 | 0,27 | 0,0031 | 1,10 |
| Effect Tests | | | | | |
| Source | Nparm | DF | Sum of Squares | F Ratio | Prob > F |
| Place{Gribskov&Solhomfjell-Finse&Vettakollen&Svalbard} | 1 | 1 | 384,43 | 43,35 | <.0001 |
| Place{Gribskov-Solhomfjell} | 1 | 1 | 48,95 | 5,52 | 0,0196 |
| Type_genus{Trametes&Microcephale&Pluteus&Naucoria&Hygrophoropsis&Cystoderma&Lycoperdon&Rhodocollybia&Calvatia&Mycena&Galerina&Amanita&Helvella&Hypholoma-Lepista&Lactarius&Laccaria&Collybia&Ramaria&Russula&Inocybe&Cortinarius&Suillus&Rhizopogon&Hydnellum&Agaricus&Tricholoma&Hydnum} | 1 | 1 | 1884,77 | 212,54 | <.0001 |
| Type_genus{Trametes&Microcephale&Pluteus&Naucoria&Hygrophoropsis&Cystoderma-Lycoperdon&Rhodocollybia&Calvatia&Mycena&Galerina&Amanita&Helvella&Hypholoma} | 1 | 1 | 176,10 | 19,86 | <.0001 |

Table S13. Results from the stepwise regression models on the $^{15}$N isotope data.

| **Response d13C** | |
|---|---|
| RSquare | 0,672 |
| RSquare Adj | 0,659 |
| Root Mean Square Error | 1,05 |

| | | | | | |
|---|---|---|---|---|---|
| Mean of Response | -24,73 | | | | |
| Observations (or Sum Wgts) | 252 | | | | |
| Analysis of Variance | | | | | |
| Source | DF | Sum of Squares | Mean Square | F Ratio | |
| Model | 9 | 548,2 | 60,9 | 55,0 | |
| Error | 242 | 268,1 | 1,1 | | Prob > F |
| C. Total | 251 | 816,4 | | <.0001 | |
| Parameter Estimates | | | | | |
| Term | %Variance | Estimate | Std Error | Prob>\|t\| | VIF |
| Intercept | -- | -25,66 | 0,26 | <.0001 | -- |
| Place{Svalbard&Finse&Gribskov-Solhomfjell&Vettakollen} | 10,3 | -0,53 | 0,07 | <.0001 | 1,1 |
| Place{Svalbard-Finse&Gribskov} | 1,4 | -0,25 | 0,09 | 0,0075 | 1,1 |
| %N | 2,8 | 0,175 | 0,045 | 0,0001 | 1,3 |
| Type_genus**{Lactarius&Russula&Helvella&Naucoria&Amanita&Tricholoma&Laccaria&Cortinarius**&*Inocybe&Galerina&Microcephale-Hydnum&Pluteus&Lycoperdon&Hydnellum&Cystoderma*&Rhizopogon&Mycena&Calvatia&Lepista&Rhodocollybia&Ramaria&**Hypholoma&Suillus&Trametes&Agaricus&Collybia&Hygrophoropsis**} | 30,6 | -1,15 | 0,09 | <.0001 | 1,7 |
| Type_genus{Lactarius&Russula&Helvella&Naucoria&Amanita&Tricholoma&Laccaria&Cortinarius-Inocybe&Galerina&Microcephale} | 3,8 | -0,58 | 0,13 | <.0001 | 1,3 |
| Type_genus{Amanita&Tricholoma-Laccaria&Cortinarius} | 1,2 | -0,49 | 0,19 | 0,0119 | 1,1 |
| Type_genus{Hydnum&Pluteus&Lycoperdon&Hydnellum&Cystoderma&Rhizopogon&Mycena&Calvatia&Lepista&Rhodocollybia&Ramaria-Hypholoma&Suillus&Trametes&Agaricus&Collybia&Hygrophoropsis} | 11,6 | -0,97 | 0,12 | <.0001 | 1,7 |
| Type_genus{Hydnum&Pluteus&Lycoperdon&Hydnellum&Cystoderma-Rhizopogon&Mycena&Calvatia&Lepista&Rhodocollybia&Ramaria} | 2,6 | -0,55 | 0,15 | 0,0003 | 1,8 |
| Type_genus{Hypholoma&Suillus-Trametes&Agaricus&Collybia&Hygrophoropsis} | 1,5 | -0,56 | 0,20 | 0,0048 | 1,1 |
| Type_genus**{Lactarius&Russula&Helvella&Naucoria&Amanita&Tricholoma&Laccaria&Cortinarius**&*Inocybe&Galerina&Microcephale-Hydnum&Pluteus&Lycoperdon&Hydnell* | 30,6 | -1,15 | 0,09 | <.0001 | 1,7 |

| | | | | | |
|---|---|---|---|---|---|
| *um&Cystoderma*&Rhizopogon&Mycena&Calvatia&Lepista&Rhodocollybia&Ramaria&**Hypholoma&Suillus&Trametes&Agaricus&Collybia&Hygrophoropsis**} | | | | | |
| Effect Tests | | | | | |
| Source | Nparm | %Variance | Sum of Squares | F Ratio | Prob > F |
| Place{Svalbard&Finse&Gribskov-Solhomfjell&Vettakollen} | 1 | 10,3 | 60,4 | 54,5 | <.0001 |
| Place{Svalbard-Finse&Gribskov} | 1 | 1,4 | 8,1 | 7,3 | 0,0075 |
| %N | 1 | 2,8 | 16,6 | 15,0 | 0,0001 |
| Type_genus{Lactarius&Russula&Helvella&Naucoria&Amanita&Tricholoma&Laccaria&Cortinarius&Inocybe&Galerina&Microcephale-Hydnum&Pluteus&Lycoperdon&Hydnellum&Cystoderma&Rhizopogon&Mycena&Calvatia&Lepista&Rhodocollybia&Ramaria&Hypholoma&Suillus&Trametes&Agaricus&Collybia&Hygrophoropsis} | 1 | 30,6 | 178,8 | 161,4 | <.0001 |
| Type_genus{Lactarius&Russula&Helvella&Naucoria&Amanita&Tricholoma&Laccaria&Cortinarius-Inocybe&Galerina&Microcephale} | 1 | 3,8 | 22,4 | 20,2 | <.0001 |
| Type_genus{Amanita&Tricholoma-Laccaria&Cortinarius} | 1 | 1,2 | 7,1 | 6,4 | 0,0119 |
| Type_genus{Hydnum&Pluteus&Lycoperdon&Hydnellum&Cystoderma&Rhizopogon&Mycena&Calvatia&Lepista&Rhodocollybia&Ramaria-Hypholoma&Suillus&Trametes&Agaricus&Collybia&Hygrophoropsis} | 1 | 11,6 | 67,6 | 61,1 | <.0001 |
| Type_genus{Hydnum&Pluteus&Lycoperdon&Hydnellum&Cystoderma-Rhizopogon&Mycena&Calvatia&Lepista&Rhodocollybia&Ramaria} | 1 | 2,6 | 15,2 | 13,7 | 0,0003 |
| Type_genus{Hypholoma&Suillus-Trametes&Agaricus&Collybia&Hygrophoropsis} | 1 | 1,5 | 9,0 | 8,1 | 0,0048 |

Table S14. Results from the stepwise regression models on the $^{13}$C isotope data.

| **GenBank Accession number** | **Genbank species name** | **Probable identification** |
|---|---|---|
| GU234118.1 | *Mycena filopes* | *Mycena metata* |
| JN021065.2 | *M.cf.purpureofusca* | *not M.purpureofusca* |
| JQ676208.1 | *M.cf.purpureofusca* | *not M. purpureofusca* |
| JF908390.1 | *M. vitilis* | *M. sanguinolenta* |
| DQ384588.1 | *M. vitilis* | *not M. vitilis.* |
| MH856158 | *M.galopus* | *M. epipterygia* |

| | | |
|---|---|---|
| GU234146 | *M. cinerella* | *M. strobilinoidea.* |
| UDB015275 | *M. leptocephala* | *not M. leptocephala* |
| JF908471 | *M. rosella* | *not M.rosella.* |
| JF908473 | *M. rosella* | *M.rosea* |
| KX449424 | *M. rosella* | *M.rosea* |
| JF908487 | *M. rosella* | *M.rosea* |
| JF908488 | *M. rosella* | *M.rosea* |
| KR673440.1 | *M. arcangeliana* | *not M.arcangeliana* |

Table S15. 14 full-length ITS sequences from GenBank that were deemed as misidentified, as they were either single sequences or multiple sequences from one source that showed less than 95% similarity with several other sequences from supposedly conspecific collections and/or fell systematically with a cluster of multiple sequences from another species.

| **Previously published datasets** | **Source for downloading** |
|---|---|
| **Davey et al. (2015)** | **NCBI (SRP006874).** |
| **Mundra et al. (2015)** | **Dryad (doi:10.5061/dryad.2343k).** |
| **Yao et al. (2013)** | **Dryad (doi:10.5061/dryad.216tp).** |
| **Lorberau et al. (2017)** | **Dryad (doi:10.5061/dryad.49dn0).** |
| **Botnen et al. (2014)** | **Dryad (doi.org/10.5061/dryad.45pv2)** |
| **Botnen et al. (2019)** | **Dryad (doi.org/10.5061/ dryad.n42dd20.)** |
| **Blaalid et al. (2012)** | **European Nucleotide Archive; accession no. SRP006836.1** |
| **Blaalid et al. (2014):** | **NCBI short read archive, accession no. SRP006836** |
| **Jarvis et al. (2015):** | **NCBI short read archive, accession no. (PRJNA253816).** |
| **New data** | **Source for downloading** |
| ***B. pubescens*** | **NCBI (archive PRJNA706760)** |
| ***A. alpinus, A.uva-ursi, B.nana, S. herbacea*.** | **Zenodo Digital identifier: 10.5281/zenodo.4545738** |

Table S16. Data sources for the 454/Illumina data.

| **Sequence** | **Organism** | **Specimen-Voucher** | **FINBOL-no.** |
|---|---|---|---|
| MW540654 | Mycena aff. polygramma | H6029504 | FIMY001-12 |
| MW540655 | Phloeomana sp. | H6029500 | FIMY005-12 |
| MW540656 | Mycena niveipes | H6030712 | FIMY016-12 |
| MW540657 | Mycena aurantiomarginata | H6032424 | FIMY020-12 |
| MW540658 | Mycena pearsoniana | H6032595 | FIMY022-12 |
| MW540659 | Mycena abramsii | H6032598 | FIMY025-12 |
| MW540660 | Mycena aff. pearsoniana | H6032605 | FIMY032-12 |
| MW540662 | Mycena galericulata | H6032610 | FIMY035-12 |
| MW540663 | Mycena galopus | H6032614 | FIMY037-12 |
| MW540664 | Mycena tintinnabulum | H6008524 | FIMY041-12 |

| | | | |
|---|---|---|---|
| MW540665 | Mycena silvae-nigrae | H6004009 | FIMY046-12 |
| MW540666 | Mycena aff. niveipes | H6033550 | FIMY047-12 |
| MW540667 | Mycena plumipes | H6033551 | FIMY052-12 |
| MW540668 | Mycena latifolia | H6033552 | FIMY053-12 |
| MW540669 | Mycena sanguinolenta | H6033553 | FIMY054-12 |
| MW540670 | Mycena cinerella | H6018219 | FIMY055-12 |
| MW540671 | Mycena leptocephala | H6029112 | FIMY064-13 |
| MW540672 | Mycena lammiensis | H6008503 | FIMY068-13 |
| MW540674 | Mycena clavicularis | H6036816 | FIMY070-13 |
| MW540675 | Mycena adscendens | H6036818 | FIMY073-13 |
| MW540678 | Mycena mucor | H6036830 | FIMY075-13 |
| MW540679 | Mycena galopus | H6036831 | FIMY077-13 |
| MW540680 | Mycena megaspora | H6036833 | FIMY081-13 |
| MW540681 | Mycena leptocephala | H6036834 | FIMY082-13 |
| MW540682 | Mycena alexandri | H6036838 | FIMY089-13 |
| MW540683 | Mycena vulgaris | H6036839 | FIMY090-13 |
| MW540686 | Mycena haematopus | H6036847 | FIMY092-13 |
| MW540687 | Mycena amicta | H6036851 | FIMY093-13 |
| MW540688 | Mycena aetites | H6036854 | FIMY097-13 |
| MW540689 | Mycena erubescens | H6036859 | FIMY098-13 |
| MW540692 | Mycena filopes | H6036864 | FIMY100-13 |
| MW540693 | Mycena septentrionalis | H6036865 | FIMY103-13 |
| MW540694 | Mycena olivaceomarginata | H6036868 | FIMY106-13 |
| MW540695 | Mycena laevigata | H6036869 | FIMY110-13 |
| MW540696 | Mycena cyanorrhiza | J24082010 | FIMY113-13 |
| MW540697 | Mycena rosea | H6008519 | FIMY118-13 |
| MW540698 | Mycena zephirus | H6038555 | FIMY119-13 |
| MW540699 | Mycena tintinnabulum | H6012651 | FIMY122-13 |
| MW540700 | Mycena aff. rubromarginata | H6018205 | FIMY123-13 |
| MW540701 | Mycena pterigena | H6038561 | FIMY124-13 |
| MW540702 | Mycena rosella | H6039057 | FIMY127-13 |
| MW540703 | Mycena polygramma | H6039058 | FIMY128-13 |
| MW540704 | Mycena tubaroides | H6039061 | FIMY129-13 |
| MW540705 | Mycena viridimarginata | H6039063 | FIMY137-13 |
| MW540706 | Mycena vitilis | H6039064 | FIMY143-13 |
| MW540708 | Mycena rubromarginata | H6039068 | FIMY151-13 |
| MW540709 | Mycena polyadelpha | H6039077 | FIMY157-13 |
| MW540710 | Mycena pura | H6039078 | FIMY162-13 |
| MW540711 | Mycena polygramma | H6039079 | FIMY164-13 |
| MW540712 | Mycena pura | H6039083 | FIMY165-13 |
| MW540713 | Mycena mirata | TUR168621 | FIMY168-13 |
| MW540714 | Mycena longiseta | TUR173933 | FIMY170-13 |
| MW540715 | Mycena simia | TUR157134 | FIMY171-13 |

| | | | |
|---|---|---|---|
| MW540716 | Mycena cf. rapiolens | TUR171435 | FIMY173-13 |
| MW540717 | Mycena picta | TUR194167 | FIMY175-13 |
| MW540718 | Mycena aff. algeriensis | TUR182370 | FIMY184-13 |
| MW540720 | Mycena rorida | TUR136996 | FIMY185-13 |
| MW540722 | Mycena maculata | TUR157124 | FIMY186-13 |
| MW540723 | Mycena tristis | TUR168636 | FIMY190-13 |
| MW540725 | Mycena rebaudengoi | F073581 | FIMY196-14 |
| MW540726 | Mycena aff. pasvikensis | H6045675 | FIMY197-14 |
| MW540727 | Mycena aff. epipterygia | H6045676 | FIMY199-14 |
| MW540728 | Mycena stylobates | F073091 | FIMY200-14 |
| MW540729 | Mycena capillaripes | F074461 | FIMY201-14 |
| MW540730 | Mycena cf. concolor | F018012 | FIMY203-14 |
| MW540731 | Mycena olivaceomarginata | F076595 | FIMY204-14 |
| MW540732 | Mycena cf. mucolor | F033378 | FIMY209-14 |
| MW540734 | Mycena cf. cineroides | F043625 | FIMY214-14 |
| MW540736 | Mycena epipterygia | H6050643 | FIMY215-14 |
| MW540737 | Mycena cf. epipterygia | H6045797 | FIMY218-14 |
| MW540738 | Mycena metata | H6057234 | FIMY224-14 |
| MW540739 | Mycena arcangeliana | TUR148293 | FIMY227-14 |
| MW540740 | Mycena pelianthina | TUR177351 | FIMY243-14 |
| MW540741 | Mycena aff. meliigena | H6057961 | FIMY244-14 |
| MW540742 | Mycena aff. alexandri | TUR181346 | |
| MW576881 | Mycena rosella | CBHHK_SolIII | |
| MW576882 | Mycena rosella | CBHHK_SolII | |
| MW576883 | Mycena galopus | CBHHK_SolI | |
| MW576884 | Mycena rosella | CBHHK_SolV | |
| MW576885 | Mycena metata | CBHHK_Sol_IV | |
| MW576886 | Mycena galopus | CBHHK_Telemark2 | |
| MW576887 | Mycena vitilis | CBHHK_100 | |
| MW576888 | Mycena galopus | CBHHK_102 | |
| MW576890 | Mycena metata | CBHHK_103 | |
| MW576891 | Mycena galopus | CBHHK_105 | |
| MW576892 | Mycena metata | CBHHK_107 | |
| MW576893 | Mycena metata | CBHHK_110 | |
| MW576894 | Mycena galopus | CBHHK_112 | |
| MW576895 | Mycena galopus | CBHHK_117 | |
| MW576896 | Mycena metata | CBHHK_118 | |
| MW576897 | Mycena metata | CBHHK_123 | |
| MW576898 | Mycena epipterygia | CBHHK_124 | |
| MW576899 | Mycena metata | CBHHK_125 | |
| MW576900 | Mycena metata | CBHHK_126 | |
| MW576901 | Mycena galopus | CBHHK_130 | |
| MW576902 | Mycena galopus | CBHHK_133 | |
| MW576903 | Mycena epipterygia | CBHHK_136 | |

| | | |
|---|---|---|
| MW576904 | Mycena epipterygia | CBHHK_139 |
| MW576905 | Mycena maculata | CBHHK_140m |
| MW576906 | Mycena metata | CBHHK_143 |
| MW576907 | Mycena vulgaris | CBHHK_146 |
| MW576908 | Mycena epipterygia | CBHHK_147 |
| MW576909 | Mycena vitilis | CBHHK_149 |
| MW576910 | Mycena metata | CBHHK_151m |
| MW576911 | Mycena galericulata | CBHHK_152 |
| MW576912 | Mycena metata | CBHHK_154m |
| MW576913 | Mycena rosella | CBHHK_155m |
| MW576914 | Mycena galericulata | CBHHK_161 |
| MW576915 | Mycena crocata | CBHHK_163 |
| MW576916 | Mycena crocata | CBHHK_164m |
| MW576917 | Mycena vitilis | CBHHK_166m |
| MW576918 | Mycena vitilis | CBHHK_167 |
| MW576919 | Mycena vitilis | CBHHK_168m |
| MW576920 | Mycena metata | CBHHK_170 |
| MW576921 | Mycena vitilis | CBHHK_171 |
| MW576922 | Mycena metata | CBHHK_172m |
| MW576923 | Mycena belliae | CBHHK_174m |
| MW576924 | Mycena sanguinolenta | CBHHK_176m |
| MW576925 | Mycena metata | CBHHK_178 |
| MW576926 | Mycena vitilis | CBHHK_179m |
| MW576927 | Mycena metata | CBHHK_182m |
| MW576928 | Mycena crocata | CBHHK_184m |
| MW576929 | Mycena metata | CBHHK_198 |
| MW576930 | Mycena metata | CBHHK_28 |
| MW576931 | Mycena vulgaris | CBHHK_31 |
| MW576932 | Mycena metata | CBHHK_32 |
| MW576933 | Mycena vitilis | CBHHK_34 |
| MW576934 | Mycena rosella | CBHHK_45 |
| MW576935 | Mycena galericulata | CBHHK_50 |
| MW576936 | Mycena galericulata | CBHHK_51 |
| MW576937 | Mycena rosella | CBHHK_53 |
| MW576938 | Mycena metata | CBHHK_61 |
| MW576939 | Mycena galopus | CBHHK_64m |
| MW576940 | Mycena rosella | CBHHK_65 |
| MW576941 | Mycena galopus | CBHHK_69 |
| MW576942 | Mycena metata | CBHHK_70 |
| MW576943 | Mycena metata | CBHHK_73 |
| MW576944 | Mycena sanguinolenta | CBHHK_80 |
| MW576945 | Mycena metata | CBHHK_81 |
| MW576946 | Mycena vitilis | CBHHK_85 |
| MW576947 | Mycena metata | CBHHK_88 |
| MW576948 | Mycena amicta | JBFRANK_9232 |

| | | |
|---|---|---|
| MT153145 | Mycena rorida | JBFRANK_9284 |
| MW576950 | Mycena capillaripes | JBFRANK_9286 |
| MW576951 | Mycena metata | CBHHK_93 |
| MW576952 | Mycena vitilis | CBHHK_94 |
| MW576953 | Mycena rosella | CBHHK_95 |
| MW576954 | Mycena epipterygia | CBHHK_96 |
| MW576955 | Mycena galopus | JBFRANK_9904 |
| MW576956 | Mycena vitilis | CBHHK_99 |
| MW576957 | Mycena cf.alexandri | CBHHK_A171 |
| AB512311 | Mycena sanguinolenta | |
| AB512312 | Mycena chlorophos | |
| AF335444 | Mycena aff.murina | |
| AY805614 | Mycena galopus | |
| MT153137 | Mycena latifolia | |
| MT153132 | Mycena filopes | |
| MT153125 | Mycena albidolilacea | |
| MT153149 | Mycena vulgaris | |
| MT153146 | Mycena rosella | |
| MT153144 | Mycena rebaudengoi | |
| MT153147 | Mycena sanguinolenta | |
| MT153142 | Mycena polygramma | |
| MT153131 | Mycena epipterygia | |
| MT153140 | Mycena metata | |
| MT153133 | Mycena galericulata | |
| MT153148 | Mycena vitilis | |
| MT153128 | Mycena belliae | |
| MT153130 | Mycena crocata | |
| MT153139 | Mycena maculata | |
| MT153136 | Mycena haematopus | |
| MT153143 | Mycena pura | |
| MT153126 | Mycena alexandri | |
| DQ384586 | Mycena cf.epipterygia | |
| DQ404392 | Mycena galericulata | |
| DQ490643 | Mycena aff.pura | |
| DQ490645 | Mycena amicta | |
| DQ494677 | Mycena plumbea | |
| EF530930 | Mycena maculata | |
| EF530939 | Mycena rubromarginata | |
| EF530946 | Mycena epipterygia.var.epipterygia | |
| EU486451 | Mycena epipterygia | |
| EU517504 | Mycena pura | |
| EU517505 | Mycena pura | |
| EU517506 | Mycena pura | |

| | |
|---|---|
| EU669223 | Mycena tenax |
| EU669224 | Mycena tenax |
| EU697245 | Mycena monticola |
| EU834204 | Mycena cf.quiniaultensis |
| EU846251 | Mycena tenax |
| EU846300 | Mycena hudsoniana |
| FJ596760 | Mycena rorida |
| FJ596761 | Mycena rorida |
| FJ596764 | Mycena sanguinolenta |
| FJ596884 | Mycena epipterygia |
| FJ596888 | Mycena semivestipes |
| FN394560 | Mycena dura |
| FN394562 | Mycena pura |
| FN394610 | Mycena pura |
| FN394614 | Mycena pearsoniana |
| FN394618 | Mycena diosma |
| GU054133 | Mycena chlorophos |
| GU062319 | Mycena galericulata |
| GU234095 | Mycena atroalboides |
| GU234111 | Mycena citrinomarginata |
| GU234112 | Mycena olivaceomarginata |
| GU234119 | Mycena olivaceomarginata |
| GU234138 | Mycena simia |
| GU234150 | Mycena citrinomarginata |
| GU234165 | Mycena pura |
| MT153141 | Mycena olivaceomarginata |
| MT153138 | Mycena leptocephala |
| HM240533 | Mycena epipterygia |
| HM240534 | Mycena galopus |
| HM240535 | Mycena pura |
| HM240536 | Mycena rubromarginata |
| HM240537 | Mycena rubromarginata |
| HQ604765 | Mycena purpureofusca |
| HQ604766 | Mycena purpureofusca |
| HQ604767 | Mycena purpureofusca |
| HQ604768 | Mycena haematopus |
| HQ604771 | Mycena epipterygia |
| HQ604772 | Mycena epipterygia |
| HQ604774 | Mycena tenerrima |
| JF340273 | Mycena galericulata |
| JF519108 | Mycena amicta |
| JF908366 | Mycena corynephora |

| | |
|---|---|
| JF908367 | Mycena corynephora |
| JF908368 | Mycena corynephora |
| JF908369 | Mycena corynephora |
| JF908370 | Mycena sanguinolenta |
| JF908371 | Mycena angusta |
| JF908372 | Mycena rhamnicola |
| JF908374 | Mycena belliae |
| JF908375 | Mycena terena |
| JF908376 | Mycena leaiana |
| JF908377 | Mycena albidolilacea |
| JF908378 | Mycena viridimarginata |
| JF908379 | Mycena pelianthina |
| JF908380 | Mycena pelianthina |
| JF908385 | Mycena cyanorhiza |
| JF908386 | Mycena pseudocorticola |
| JF908387 | Mycena pseudocorticola |
| JF908388 | Mycena supina |
| JF908389 | Mycena supina |
| JF908391 | Mycena renati |
| JF908392 | Mycena strobilinoidea |
| JF908393 | Mycena strobilinoidea |
| JF908394 | Mycena amicta |
| JF908400 | Mycena abramsii |
| JF908401 | Mycena arcangeliana |
| JF908402 | Mycena arcangeliana |
| JF908403 | Mycena pura |
| JF908407 | Mycena megaspora |
| JF908410 | Mycena filopes |
| JF908415 | Mycena citrinomarginata |
| JF908416 | Mycena citrinomarginata |
| JF908417 | Mycena diosma |
| JF908420 | Mycena adscendens |
| JF908421 | Mycena romagnesiana |
| JF908422 | Mycena romagnesiana |
| JF908423 | Mycena meliigena |
| JF908424 | Mycena algeriensis |
| JF908425 | Mycena algeriensis |
| JF908426 | Mycena alnetorum |
| JF908427 | Mycena calceata |
| JF908428 | Mycena meliigena |
| JF908429 | Mycena meliigena |
| JF908430 | Mycena rubromarginata |
| JF908433 | Mycena polygramma |
| JF908434 | Mycena polygramma |
| JF908435 | Mycena vulgaris |

| | |
|---|---|
| JF908439 | Mycena stylobates |
| JF908440 | Mycena strobilicola |
| JF908441 | Mycena galericulata |
| JF908442 | Mycena galericulata |
| JF908443 | Mycena capillaris |
| JF908449 | Mycena pura |
| JF908450 | Mycena pura |
| JF908451 | Mycena pura |
| JF908452 | Mycena silvaenigrae |
| JF908453 | Mycena silvaenigrae |
| JF908455 | Mycena niveipes |
| JF908456 | Mycena polyadelpha |
| JF908457 | Mycena subcana |
| JF908458 | Mycena epipterygia |
| JF908460 | Mycena epipterygia |
| JF908461 | Mycena zephirus |
| JF908462 | Mycena zephirus |
| JF908463 | Mycena pilosella |
| JF908465 | Mycena valida |
| JF908466 | Mycena clavicularis |
| JF908467 | Mycena clavicularis |
| JF908469 | Mycena seynesii |
| JF908470 | Mycena seynesii |
| JF908472 | Mycena pura |
| JF908473 | Mycena rosea |
| JF908475 | Mycena cupressina |
| JF908477 | Mycena rebaudengoi |
| JF908478 | Mycena juniperina |
| JF908479 | Mycena aurantiomarginata |
| JF908480 | Mycena albidorosea |
| JF908481 | Mycena graminicola |
| JF908483 | Mycena thymicola |
| JF908484 | Mycena galopus |
| JF908485 | Mycena latifolia |
| JF908486 | Mycena rhamnicola |
| JF908487 | Mycena rosea |
| JF908488 | Mycena rosea |
| JF908490 | Mycena albidoaquosa |
| JF908491 | Mycena pachyderma |
| JF908492 | Mycena crocata |
| JN182198 | Mycena pearsoniana |
| JN182199 | Mycena pearsoniana |
| JN182200 | Mycena pearsoniana |
| JN182201 | Mycena pearsoniana |

| | |
|---|---|
| JN182202 | Mycena pura |
| JN198391 | Mycena plumbea |
| JQ358808 | Mycena laevigata |
| JQ358809 | Mycena purpureofusca |
| JQ358810 | Mycena rubromarginata |
| JQ364945 | Mycena purpureofusca |
| JQ926166 | Mycena galopus |
| JX297424 | Mycena plumipes |
| JX297425 | Mycena plumipes |
| JX297426 | Mycena plumipes |
| JX297427 | Mycena plumipes |
| JX310425 | Mycena monticola |
| JX434650 | Mycena cf.sanguinolenta |
| JX481737 | Mycena deeptha |
| KC581347 | Mycena pura |
| KC876328 | Mycena silvaenigrae |
| KC965695 | Mycena urania |
| KF007948 | Mycena pura |
| KF010856 | Mycena chlorophos |
| KF359604 | Mycena silvaenigrae |
| KF537247 | Mycena sinar |
| KF537248 | Mycena cahaya |
| KF537249 | Mycena sinar.var.tangkaisinar |
| KF537250 | Mycena seminau |
| KF537251 | Mycena sinar.var.tangkaisinar |
| KF537252 | Mycena seminau |
| KF668293 | Mycena galopus |
| KF668294 | Mycena sanguinolenta |
| KF668310 | Mycena haematopoda |
| KF692075 | Mycena haematopus |
| KF913022 | Mycena pura |
| KF913023 | Mycena pura |
| KJ093496 | Mycena aff.murina |
| KJ144653 | Mycena aff.pura |
| KJ206965 | Mycena chlorophos |
| KJ206966 | Mycena noctilucens |
| KJ206967 | Mycena chlorophos |
| KJ206968 | Mycena chlorophos |
| KJ206969 | Mycena chlorophos |
| KJ206970 | Mycena chlorophos |
| KJ206971 | Mycena chlorophos |
| KJ206972 | Mycena chlorophos |
| KJ206973 | Mycena chlorophos |

| | |
|---|---|
| KJ206974 | Mycena chlorophos |
| KJ206975 | Mycena illuminans |
| KJ206976 | Mycena illuminans |
| KJ206980 | Mycena illuminans |
| KJ206983 | Mycena chlorophos |
| KJ206985 | Mycena chlorophos |
| KJ206986 | Mycena chlorophos |
| KJ609168 | Mycena chlorophos |
| KJ705175 | Mycena filopes |
| KJ705176 | Mycena leaiana.var.leaiana |
| KJ705177 | Mycena vulgaris |
| KJ705178 | Mycena galericulata |
| KJ705179 | Mycena robusta |
| KJ705180 | Mycena citrinomarginata |
| KJ705181 | Mycena haematopus |
| KJ705182 | Mycena maculata |
| KJ705183 | Mycena maculata |
| KJ705184 | Mycena pura |
| KJ705186 | Mycena pura |
| KJ705187 | Mycena pura |
| KJ705188 | Mycena amicta |
| KJ713981 | Mycena pura |
| KJ831841 | Mycena chlorophos |
| KM085362 | Mycena galericulata |
| KM085398 | Mycena alnetorum |
| KM282283 | Mycena galericulata |
| KP406534 | Mycena epipterygia |
| KP454009 | Mycena rubromarginata |
| KP454034 | Mycena epipterygia.var.lignicola |
| KR673438 | Mycena arcangeliana |
| KR673481 | Mycena abramsii |
| KR673599 | Mycena galericulata |
| KR673702 | Mycena haematopus |
| KT222190 | Mycena dura |
| KT695316 | Mycena haematopus |
| KT900140 | Mycena adscendens |
| KT900141 | Mycena adscendens |
| KT900142 | Mycena adscendens |
| KT900143 | Mycena adscendens |
| KT900144 | Mycena alexandri |
| KT900145 | Mycena alexandri |
| KT900146 | Mycena cinerella_Aronsen051014 |

| | |
|---|---|
| KU295552 | Mycena alnetorum |
| KU516418 | Mycena galopus |
| KU516419 | Mycena galopus |
| KU516420 | Mycena galopus |
| KU516421 | Mycena galopus |
| KU518323 | Mycena haematopus |
| KU861555 | Mycena pasvikensis_AAronsen11111 |
| KU861556 | Mycena pasvikensis_AAronsen8612 |
| KU861557 | Mycena pasvikensis |
| KU861558 | Mycena pasvikensis |
| KU861559 | Mycena pasvikensis_AAronsen2814 |
| KU861565 | Mycena mucor_AAronsen5140914 |
| KU861566 | Mycena mucor_AAronsen7051113 |
| KU861567 | Mycena polyadelpha(AAronsen8261013) |
| KX010907 | Mycena deformis |
| KX010908 | Mycena globulispora |
| KX010909 | Mycena oculisnymphae |
| KX010910 | Mycena oculisnymphae |
| KX058336 | Mycena plumbea |
| KX236103 | Mycena maculata |
| KX449424 | Mycena rosea |
| KX449443 | Mycena inclinata |
| KX513844 | Mycena bulliformis |
| KY352524 | Mycena abramsii |
| KY681454 | Mycena zephirus |
| KY744173 | Mycena albiceps |
| KY950446 | Mycena galericulata |
| LC314114 | Mycena abramsii |
| LC373247 | Mycena diosma |
| MF161203 | Mycena haematopus |
| MF417759 | Mycena citricolor |
| MF417760 | Mycena citricolor |
| MF417761 | Mycena citricolor |
| MF417762 | Mycena citricolor |

| | |
|---|---|
| MF417763 | Mycena citricolor |
| MF686517 | Mycena haematopus |
| MF686520 | Mycena leaiana |
| MF773619 | Mycena haematopus |
| MF908474 | Mycena maculata |
| MF926553 | Mycena cf.cinerella |
| MF926554 | Mycena cf.cinerella |
| MF943121 | Mycena purpureofusca |
| MF955190 | Mycena cf.pura |
| MF955191 | Mycena cf.pura |
| MF955192 | Mycena cf.pura |
| MF993026 | Mycena indigotica |
| MG654739 | Mycena citrinomarginata |
| MG654740 | Mycena purpureofusca |
| MG654741 | Mycena purpureofusca |
| MG654742 | Mycena purpureofusca |
| MG654743 | Mycena strobilinoidea |
| MG654744 | Mycena strobilinoidea |
| MG719609 | Mycena haematopus |
| MG719614 | Mycena filopes |
| MG719769 | Mycena plumbea |
| MG738261 | Mycena haematopus |
| MG748570 | Mycena niveipes |
| MG926696 | Mycena globulispora |
| MG926697 | Mycena globulispora |
| MG969987 | Mycena citrinomarginata |
| MG969988 | Mycena citrinomarginata |
| MH063433 | Mycena indigotica |
| MH136830 | Mycena alphitophora |
| MH136831 | Mycena alphitophora |
| MH136832 | Mycena corynephora |
| MH136833 | Mycena corynephora |
| MH136834 | Mycena corynephora |
| MH142010 | Mycena haematopus |
| MH142012 | Mycena haematopus |
| MH145355 | Mycena amicta |
| MH380201 | Mycena filopes |
| MH396626 | Mycena abramsii |
| MH396627 | Mycena abramsii |
| MH396628 | Mycena abramsii |
| MH396629 | Mycena abramsii |
| MH396630 | Mycena epipterygia |
| MH396631 | Mycena epipterygia |
| MH396632 | Mycena epipterygia |
| MH396633 | Mycena epipterygia |

| | |
|---|---|
| MH396634 | Mycena filopes |
| MH396635 | Mycena filopes |
| MH396636 | Mycena metata |
| MH396637 | Mycena metata |
| MH414547 | Mycena aff.holoporphyra |
| MH414551 | Mycena breviseta |
| MH414554 | Mycena aff.discobasis |
| MH414555 | Mycena aff.discobasis |
| MH414556 | Mycena discogena |
| MH414557 | Mycena lasiopus |
| MH414558 | Mycena lasiopus |
| MH414562 | Filoboletus_pallescens |
| MH414563 | Filoboletus_pallescens |
| MH718251 | Mycena polygramma |
| MH856225 | Mycena aetites |
| MH856226 | Mycena aetites |
| MH856227 | Mycena olivaceomarginata |
| MH856228 | Mycena olivaceomarginata |
| MH856229 | Mycena olivaceomarginata |
| MH856231 | Mycena maculata |
| MH856232 | Mycena maculata |
| MH856233 | Mycena maculata |
| MH856234 | Mycena maculata |
| MH856235 | Mycena polygramma |
| MH856236 | Mycena polygramma |
| MH856237 | Mycena polygramma |
| MH856238 | Mycena polygramma |
| MH856239 | Mycena polygramma |
| MH856240 | Mycena vulgaris |
| MH856332 | Mycena capillaripes |
| MH856333 | Mycena cinerella |
| MH856334 | Mycena olivascens |
| MH856335 | Mycena rubromarginata |
| MH856336 | Mycena rubromarginata |
| MH856337 | Mycena rubromarginata |
| MH856338 | Mycena rubromarginata |
| MH856339 | Mycena zephirus |
| MH856340 | Mycena zephirus |
| MH856341 | Mycena zephirus |
| MH856655 | Mycena amicta |
| MH856656 | Mycena citrinomarginata |
| MH856657 | Mycena |

| | |
|---|---|
| | aurantiomarginata |
| MH856658 | Mycena renati |
| MH856661 | Mycena flosnivium |
| MH856662 | Mycena sanguinolenta |
| MH856663 | Mycena xantholeuca |
| MH856664 | Mycena xantholeuca |
| MH857183 | Mycena amicta |
| MH857184 | Mycena amicta |
| MH857186 | Mycena flosnivium |
| MH857195 | Mycena citrinomarginata |
| MH857196 | Mycena citrinomarginata |
| MH857197 | Mycena citrinomarginata |
| MH857198 | Mycena citrinomarginata |
| MH857462 | Mycena alcalina |
| MH857694 | Mycena citricolor |
| MH861224 | Mycena pura.var.pura |
| MH979290 | Mycena leaiana |
| MK169369 | Mycena epipterygia.var.lignicola |
| MK169370 | Mycena overholtsii |
| MK290379 | Mycena pura |
| MK307839 | Mycena clavicularis |
| MK348517 | Mycena cf.filopes |
| MK351700 | Mycena galericulata |
| MK371751 | Mycena strobilinoidea |
| MK474930 | Mycena xantholeuca |
| MK474933 | Mycena xantholeuca |
| MK478466 | Mycena tenuispinosa |
| MK532829 | Mycena inclinata |
| MK532830 | Mycena niveipes |
| MK532831 | Mycena pura |
| MK532832 | Mycena pura |
| UDB001611 | Mycena septentrionalis |
| UDB011532 | Mycena rosea |
| UDB011648 | Mycena renati |
| UDB011668 | Mycena inclinata |
| UDB011702 | Mycena sanguinolenta |
| UDB011703 | Mycena vulgaris |
| UDB011771 | Mycena plumipes |
| UDB011809 | Mycena strobilinoidea |
| UDB011884 | Mycena epipterygia |
| UDB015405 | Mycena megaspora |
| UDB015412 | Mycena metata |
| UDB015432 | Mycena polygramma |
| UDB015495 | Mycena galericulata |

| | |
|---|---|
| UDB015861 | Mycena rosella |
| UDB016249 | Mycena polyadelpha |
| UDB016258 | Mycena mirata |
| UDB018159 | Mycena stipata |
| UDB018200 | Mycena zephirus |
| UDB019511 | Mycena laevigata |
| UDB019514 | Mycena capillaripes |
| UDB019554 | Mycena meliigena |
| UDB023691 | Mycena laevigata |
| UDB034828 | Mycena leptocephala |
| UDB037953 | Mycena maculata |

Table S17. GenBank Accession numbers for the 576 sequences in the Mycena database. Voucher information for newly sequenced specimens in the 3rd column. FinBOL-numbers (4th column) are listed for those specimens sequenced for this project as part of the FinBOL (Finnish Barcode of Life)-project under the BOLD programme (Ratnasingham and Hebert, 2007).

**Host species**

*Bistorta vivipara*, *Salix polaris* and *Dryas octopetala* are all commonly found pioneer species in the Arctic and Alpine tundras, and known to host other root-associated fungi (Hesselman, 1900; Elven et al., 2005; Bjørbækmo et al., 2010; Botnen et al., 2014). *Dryas octopetala* and *S. polaris* are woody dwarf shrubs which may produce extensive root systems, including subterraneaous runners; *B. vivipara* is a herbaceous perennial with a less extensive root system which mainly reproduces asexually by bulbils.

As an ericoid plant of the High Arctic, *Cassiope tetragona* has traditionally been believed to form ericaceous mycorrhiza (ErM) ((Strelkova, 1956; Michelsen et al., 1996)). However, (Lorberau et al., 2017) retrieved very few ErM sequences and instead found the traditionally saprotrophic genera *Clavaria* and *Mycena* and the possibly endophytic/mycorrhizal, *Sebacina* to be dominant.

*Arctostaphylos alpina* and *A. uva-ursi* are Subarctic-Alpine ericaceous plants that form arbutoid mycorrhiza where the invading hyphae form a mantle and Hartig net, but also penetrate into the outer epidermal cells of the roots (Smith and Read, 2008; Kühdorf et al., 2014).

*Salix herbacea* and *Betula nana* are ectomycorrhizal woody dwarf shrubs found on alpine and Arctic/Subarctic tundra (Mühlmann, 2008; Deslippe et al., 2011). *Pinus sylvestris* and *Betula pubescens* are both ectomycorrhizal tree hosts with a wide geographic range from boreal, temperate to subtropical areas. and which may grow up to >20 meters in height at maturity (Jonsson et al., 1999; Priha et al., 1999; Seppänen et al., 2007).

**Extended collection sites info**

The *A. uva-ursi* roots for the "altitude" study (and 9 additional *P.sylvestris* samples in addition to those from (Jarvis et al., 2015) in this study) were collected in the Invereshie-Inshriach National Nature Reserve in the north-west of the Cairngorm National Park in Scotland (Fig. 1S). At the ancient suppressed tree-line (450–500 masl), a semi-closed canopy of >5 m tall Scots pine (*Pinus sylvestris)* gives way to *Calluna-Arctostaphylos* subalpine heath. There are scattered Scots pine trees in the heath decreasing in size from the tree-line to ca. 650 masl. Bearberry extends up to 800-850masl.

Nine transects were positioned on south-west facing slopes of three adjacent mountains (Fig. 2S). Bearberry sampling locations were situated every 50 m in altitude, with transects separated by >100 m laterally. There were seven sampling points on each transect. Sample sites were located using an altimeter (Suunto vector, Vantaa, Finland) and GPS (Garmin eTrex, Southampton, UK). The lowest sampling location was within the upper limit of woodland with trees ≥5 m tall, where Scots pine roots were sampled separately.
At each sampling location, five Bearberry plants or Scots pine trees were selected within ±5 m of the target altitude (±15 m for five locations where insufficient plants were found), and 10 m distance of the determined sampling location in any direction. No other ECM hosts occurred along the transects. Fine roots were traced from main stems and three samples were collected from each plant, each containing >100 root tips. Samples were pooled at each sampling location and stored at –20 °C within 12 hours. Sampling was conducted June-July 2011.

*Betula pubescens* roots were collected at the NatureReserve at Corrimony in north-west Scotland (57°19′N 4° 43′ W). The trees are regenerating saplings at a maximum 1 m in height, growing on moorland within heather-dominated vegetation on a site heavily browsed by sheep and deer. Roots samples (supporting 100-200 ECM tips) were taken from the birches by direct tracing of fine roots from the main laterals. Roots from 5 trees from within a block were pooled to give one single sample.
A biogeography study incorporated roots sampled from ten *Arctous alpinus* sites, sixteen *Arctostaphylos uva-ursi* sites, seven *Betula nana* and seven *Salix herbacea* sites over 23 geographically distinct areas in Scotland (Hesling and Taylor 2013). These incorporated North, South, East and West extremes of the Scottish mainland Highlands, and sites on the islands of Skye and Hoy (Fig S1). Habitats included sub alpine and low alpine dwarf shrub heath, blanket bog and montane scrub. Sampling in 2010 involved collecting root samples from three positions around 10 plants within a defined sampling area which were pooled to give a single sample (Hesling & Taylor 2013). In 2011 & 2012, this was increased to 15 plants, with exceptions due to lack of suitable plants at sites: *B. nana* at Dundreggan = 8 plants, *A. alpina* Isle of Skye = 5 plants and *A. uva-ursi* Foinaven = 5 plants.
The remaining 68 *A. uva-ursi* roots for the "altitude" study (and 9 additional *P.sylvestris* samples in addition to those from (Jarvis et al., 2015) in this study) were collected June-July 2011 in the Invereshie-Inshriach National Nature Reserve in the north-west of the Cairngorm National Park in Scotland (Figs. S1, S2) at 9 transects in elevations from 450-850 masl on a *Calluna-Arctostaphylos* subalpine heath with scattered Scots pine trees up until the tree limit at ~650 masl. This was in close proximity to the mountaineous *P.sylvestris* forest studied in (Jarvis et al., 2015)

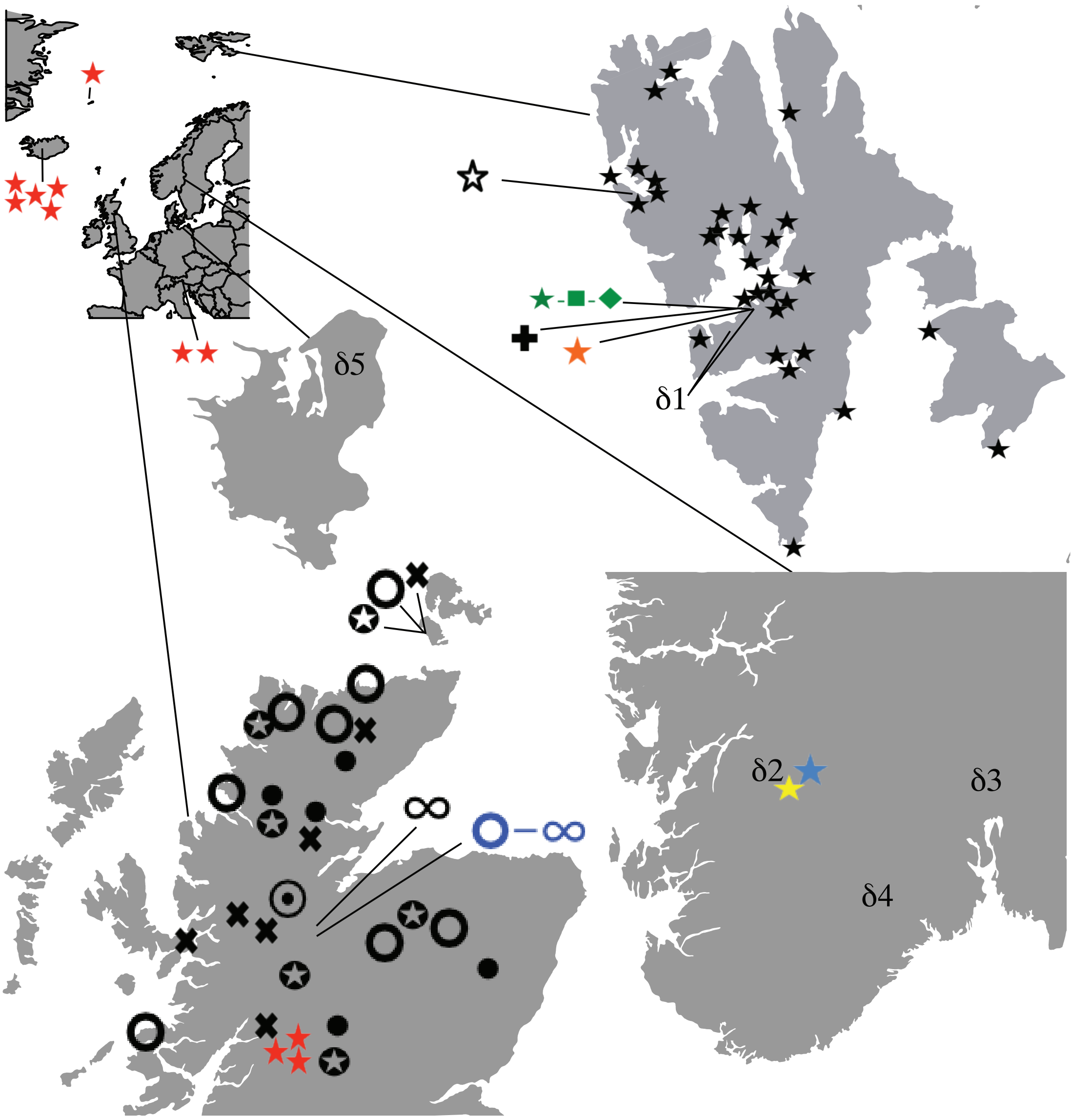

Fig.S1. Map of areas for sample origin. ★ = *Bistorta vivipara* (Blaalid et al., 2012), ★ = *Bistorta vivipara* (Yao et al., 2013). ★ = *Bistorta vivipara* (Blaalid et al 2014), ☆ = *Bistorta vivipara* Davey et al. (2015). ★-■-◆ = *Bistorta vivipara, Dryas octopetala, Salix polaris* (Botnen et al., 2014). ★ = *Bistorta vivipara* (Mundra et al., 2015), ★ = *Bistorta vivipara* (Botnen et al., 2019). □ = *Cassiope tetragona* (Lorberau et al., 2017), **O-∞** = *Arctostaphylos uva-ursi+ Pinus silvestris* (altitude study site), **O** = *Arctostaphylos uva-ursi*. □ = *Arctostaphylos alpine*. • =*Betula nana*, ⊙ = *Betula pendula*, □ = *Salix herbacea*, **∞=** *Pinus silvestris* (Jarvis et al 2015).

**d** represents places for samples for stable isotopic analyses: **d**1= Svalbard (South Isfjorden shores): Arctic tundra: Colesdalen 78°06'27.1 N 15°09'14.4E, Blomsterdalen 78°17'50.5N 17°05'15.1E, Bjørndalen 78°12'58.0N 15°19'09.6E, and Bolterdalen 78°08'23.7N 15°56'50.9E) **d**2= Finse (Mainland Norway, subarctic tundra, 60°34'53.1N, 7°29'04.1E), **d**3= Vettakollåsen (southeast Norway, mixed spruce/pine forest, 59°58'41.3N E10°42'31.9E), **d**4= Solhomfjell (southwest Norway, mixed spruce/pine forest, 58°59'10.2N 8°49'24.8E), **d**5= Gribskov (Sjælland, Denmark, beech-dominated broadleaf forest, 55°59'18.6N, 12°18'11.3E). Note that each symbol represents one sampling site from which multiple individual samples may have been collected.

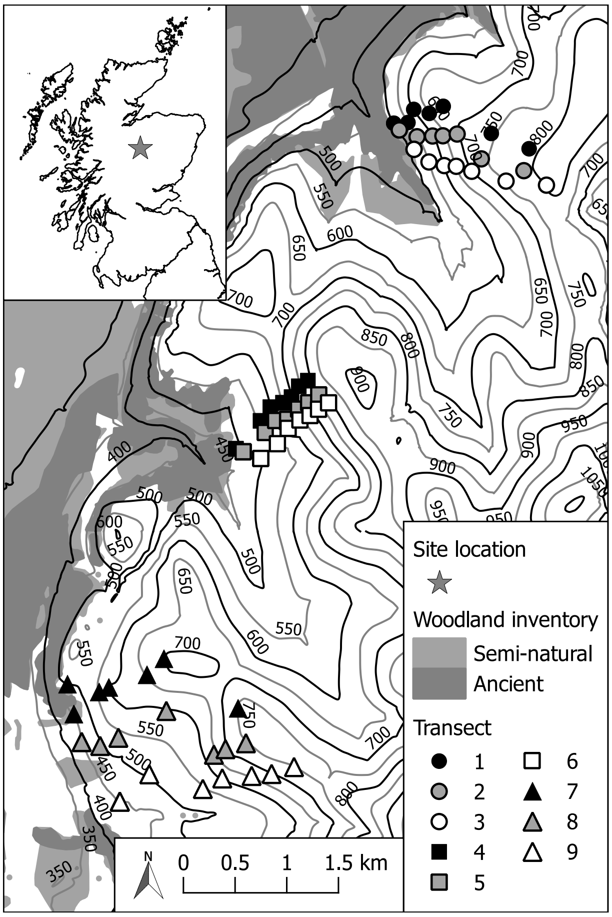

Fig.S2. Detailed map of the O-∞ = *Arctostaphylos uva-ursi/ Pinus silvestris* altitude study site in the Invereshie-Inshriach National Nature Reserve in the north-west of the Cairngorm National Park in Scotland (altitude in metres).

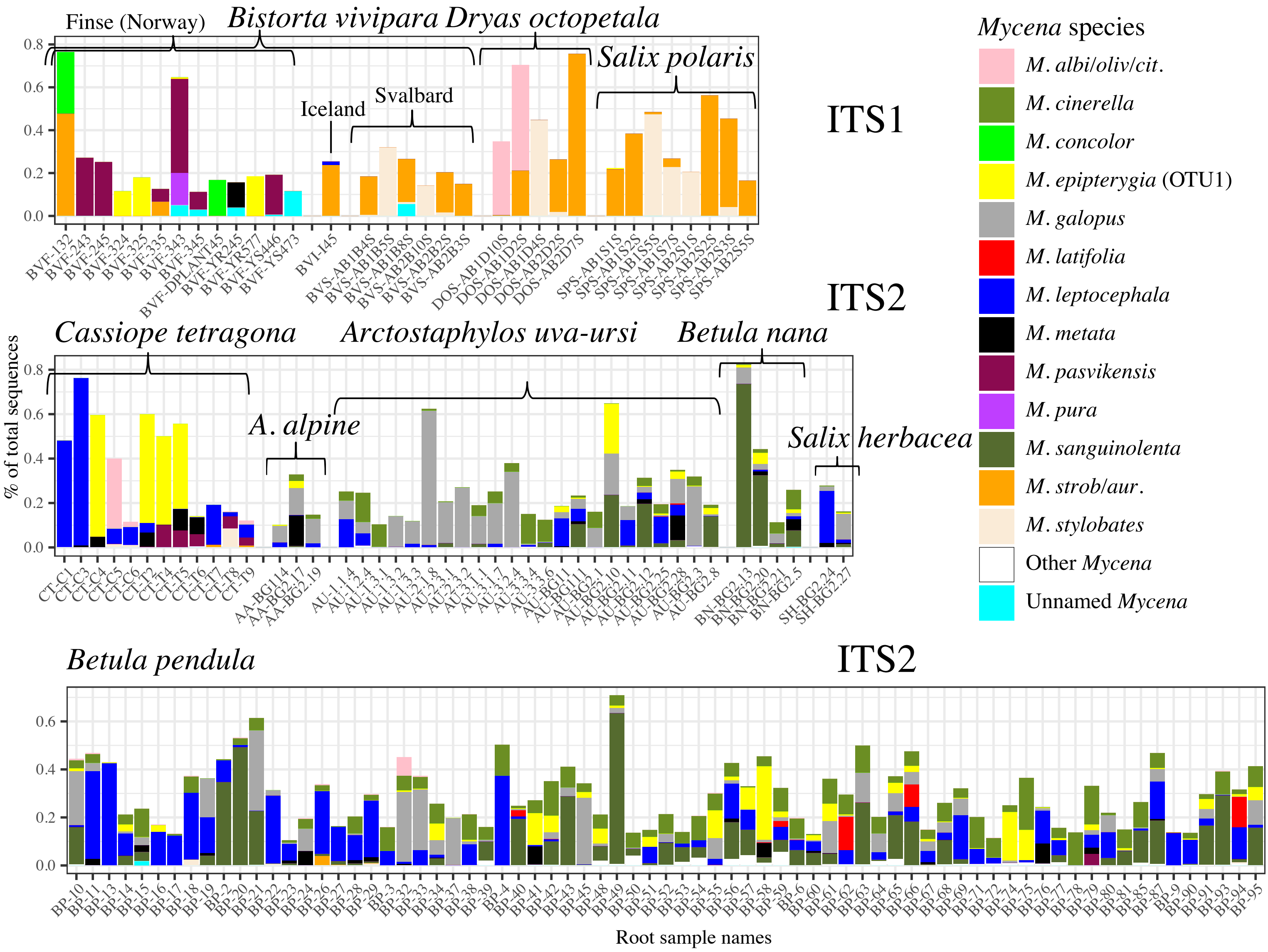

Fig.S3. Individual samples with a *Mycena* infection load (sequence content) of >10%. For *B.vivipara* from Finse (BVF_), the samples beginning with YR/YS are from Yao et al. (2013), the DPLANT45 from Blaalid et al. (2012), and the rest (BVF_[number]) from Davey et al. (2015). All *B. vivipara* from Svalbard with high *Mycena* infection rates originated from the study of Botnen et al. (2014) that also provided all *D. octopetala* + *S. polaris* samples. Note the much greater interspecific similarities in the three sets of Svalbard ITS1 samples than conspecific ditto between the *B. vivipara* from Svalbard and those from Finse. Also note the much greater similarity between *C. tetragona* from Svalbard and all the othe ITS2-based samples from Scotland than between *C. tetragona* and the other Svalbard ITS1-based samples, likely reflecting the known ITS1 reverse primer (ITS2R) bias against *Mycena*.

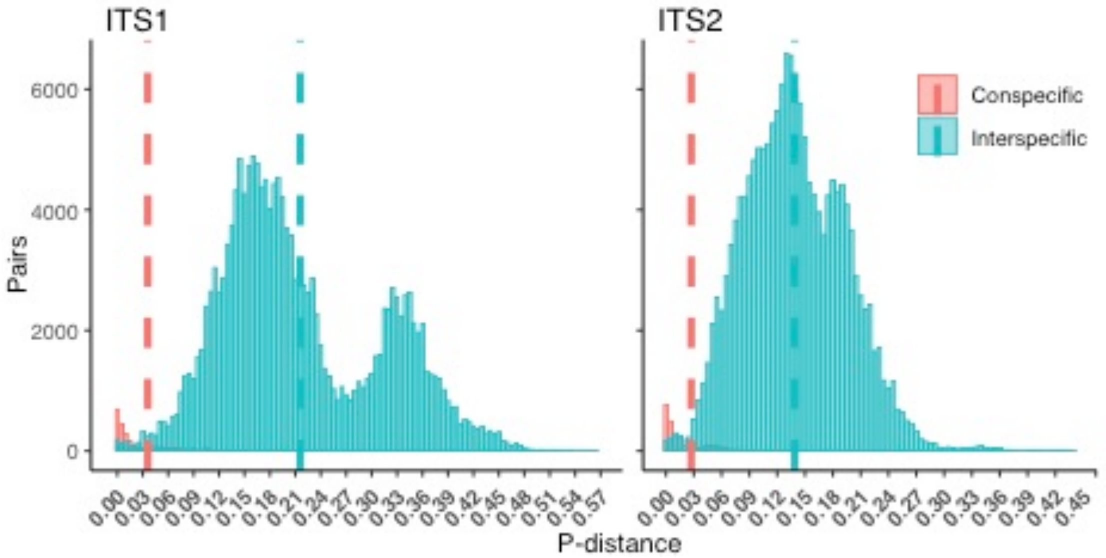

Fig.S4. Intra- and interspecific ITS1- and ITS2 distances in the 589 *Mycena* sequences that was used as a database for better species identification of *Mycena* OTUs/ASVs from HTP data sets. Note the slightly higher intraspecific variation in ITS1 (0.036) than in ITS2 (0.027), and the want of a barcoding window caused by the overlapping intra- and interspecific variation.

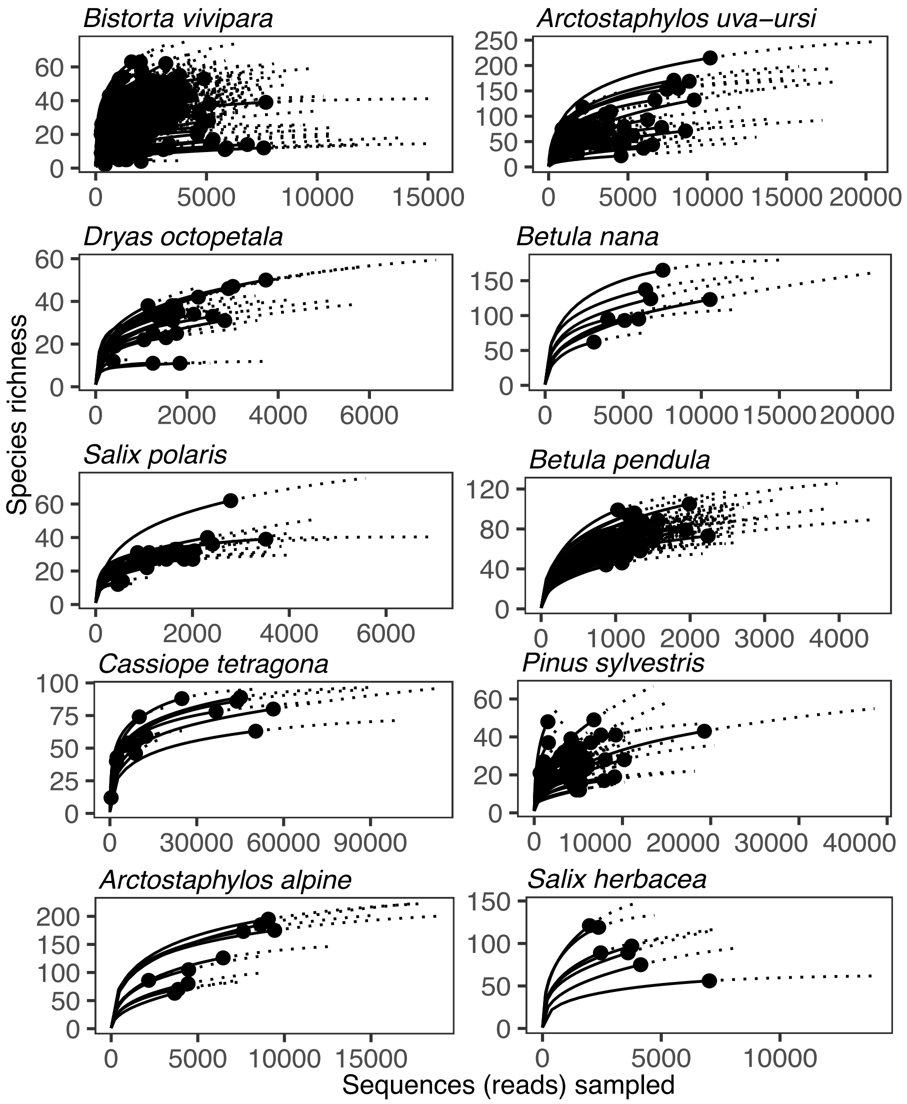

Fig.S5. iNEXT sample-based rarefaction curves with extrapolation (Hill numbers) on observed species richness. Black curves denote actual sampling (sequencing), black dots the depth limits, and dashed lines extrapolation from that up to theoretical complete coverage based on each sample.
